# Biosurfactant production under cold-saline stress is a strain specific trait independent from phylogeny and biosynthetic gene content

**DOI:** 10.64898/2026.10.19.760052

**Authors:** Habib Ullah, Zhuang Han, Maggie C. Y. Lau Vetter

## Abstract

Microbial genomes code for far more biosynthetic potential than a single cell typically expresses, but the environmental and cellular controls that determine whether the latent capacity is realised remain poorly understood. Biosurfactants, surface-active metabolites proposed to aid survival in cold and saline habitats, exemplify this gap: capacity is often inferred from genomes or detected chemically, but is rarely linked to the specific conditions that stimulate its production in the same organisms. Here we combine whole-genome sequencing, untargeted LC-MS/MS metabolomics, and quantitative surface-tension assays across 25 bacterial isolates isolated from the cold, hypersaline Qaidam Basin (a terrestrial analogue for ancient Mars) and contrasting marine environments. We show that biosurfactant production is not a fixed lineage trait but a regulated, strain-specific response that emerges under combined cold and salt stress. This response is structured into three tiers: genomic capacity defines the accessible chemical space; the genomic complement of pathway-regulator genes tracks responsiveness; strain-level physiological state determines realised output. Regulatory-gene content, rather than biosynthetic-cluster count, predicted surfactant production across strains. Production was amplified by low temperature and high salinity and showed little phylogenetic signal, with near-clonal strains differing in their stress-response. These findings reshape biochemical adaptation in extreme environments as a problem of conditional activation, not only of genomic potential, but of cellular systems that permit its expression, which requires looking beyond biosynthetic-gene content to the cellular factors that determine whether that potential is expressed.

## Introduction

Microbial genomes routinely reveal far more biosynthetic gene clusters (BGCs) than the set of specialized metabolites observed in any single culture condition. Genome mining commonly identifies tens of BGCs per genome, yet a large fraction of these remain transcriptionally silent in standard laboratory culture, creating a persistent gap between the inferred biosynthetic potential and the metabolites an organism actually produces (1,2). These BGCs are used to prioritize strains for natural-product discovery, to predict their ecological function and to interpret biochemical traces, all of which become uncertain when biosynthetic capacity prediction fails to meet the real production. This gap requires understanding not only the genome capacity but also identifying which external cues stimulate expression and which internal cellular factors gate it, which determine whether, when, and to what extent specific pathways are expressed. This is a central unresolved challenge in microbial natural-product research. The challenge is particularly acute in extreme environments, where the stress-responsive secondary metabolism may be an adaptive trait, and where the relevant environmental triggers are difficult to produce invitro. Studies that combine genomic prediction, chemical detection, and physiologically relevant stimuli in the same isolates are still rare. This limits our ability to predict which genomic BGC will be activated in nature.

Biosurfactants are a useful test case for this problem. These are surface-active amphiphiles lower interfacial tension and can contribute to maintenance stability, cryoprotection and desiccation resistance, access to hydrophobic substrate and biofilm dynamics. (3,4). These functions possibly improve survival under cold, saline, or desiccating conditions. Among biosurfactants, cyclic lipopeptides such as surfactin and lichenysin are produced non-ribosomally by *Bacillus* and related genera. These are highly potent biosurfactants and have widespread BGCs across these lineages (5,6). Critically, lipopeptide production is known to be condition dependent in model strains, ComA and other response regulator activate synthetase operon, while DegU, Spo0A and related systems modulate expression according to nutrient and stress signals (7,8). This combination of widespread genomic prevalence defined regulatory connections in model systems, and possible adaptive function makes lipopeptide biosurfactants an ideal model for asking whether this biosynthetic potential is realised as biosurfactant production under ecologically relevant stress. Yet many surveys infer biosurfactant capacity from genomes alone or from chemical detection without controlling physiological conditions, and no study is available that bring genomics, chemistry and controlled stress physiology together for the same isolates. As a result, a fundamental question remains open: for microorganisms naturally inhabiting extreme environments, is biosurfactant production a fixed property of certain lineages, a stochastic accident of cultivation, or a response actively regulated by the specific stresses those organisms encounter?

The Qaidam Basin on the northern Tibetan Plateau is a hyperarid, cold, and hypersaline endorheic basin offers an exceptional natural laboratory for addressing this question. Its microbial inhabitants experience the simultaneous stresses of a wide temperature range, fluctuating hypersalinity, intense ultraviolet radiation, and extreme desiccation, and the basin is widely recognised as one of Earth’s closest terrestrial analogues for ancient Martian lacustrine environments (9,10). Isolates from its sediments carry BGCs associated with lipopeptide production, but whether they express this capacity under the basin’s characteristic cold-saline conditions, and what determines variation in expression among closely related strains remain unknown. Because the dominant cultivable lineages of such environments often include closely related strains, the basin provides an opportunity to ask whether stress-responsive biosurfactant production is conserved phylogenetically or it varies at the strains level. Answering this is a prerequisite for predicting which of the many BGCs carried by extremophiles are actually deployed in situ, and thus for using genomic data to guide both bioprospecting and the search for metabolic biosignatures.

Here we address these questions by integrating whole-genome sequencing, untargeted liquid chromatography–tandem mass spectrometry (LC-MS/MS) metabolomics, and quantitative biosurfactant physiology across isolates from the Qaidam Basin and for comparison, isolates from contrasting deep-sea (marine) environments. By aligning genomic biosynthetic potential, the metabolites actually produced, and the surface-active phenotype expressed under controlled cold and salinity stress, we resolve biosurfactant production into three separable tiers of control. We find that production is a regulated, strain-specific and most strongly induced by the combination of low temperature and high salinity rather than being a fixed lineage trait: genomic capacity sets the outer boundary, the genomic complement of pathway-regulator genes tracks responsiveness, strain-level physiological state determines the realised output, and that this strain-specific capacity is distributed across multiple genera achieved through chemically distinct surfactant molecules, is enhanced under the combination of cold and salinity, and is separated from phylogeny even between strains with effectively identical genomes. By resolving these tiers in the same isolates, this study provide an empirical basis for a shift in how genomic potential is read in extreme environments: from cataloguing biosynthetic potential towards identifying the genomic and cellular factors that determine whether that potential is ever realised.

## Materials and Methods

### Study site, sampling, and strain collection

Microbial strains were isolated from hypersaline sediments collected from the Qaidam Basin, located on the northern Tibetan Plateau (Qinghai Province, China). The Qaidam Basin is a large endorheic basin characterised by hyperaridity (annual precipitation < 50 mm), extreme seasonal temperature fluctuations (winter minima ∼ −18 °C, summer maxima ∼ +16 °C), intense UV radiation, and dynamic salinity gradients generated by high evaporation rates (> 3,000 mm yr⁻¹), making it one of Earth’s closest terrestrial analogues to ancient Martian lacustrine environments (11–13). A set of deep-sea isolates was included to provide an ecological contrast. Full details of sample collection, field site, and initial isolation are described in Ullah et al. (under review); briefly, samples were serially diluted (10⁻¹ – 10⁻⁶) in sterile saline (20 g L⁻¹ NaCl, pH 7.2) and plated on three media, R2A agar (14), Luria–Bertani (LB) agar, and nutrient agar (NA) to maximize taxonomic recovery. Plates were incubated aerobically at 10 °C and 15 °C for 7 to 11 days. Morphologically distinct colonies were purified by repeated streaking, and pure cultures were preserved in 20% glycerol at −80 °C.

### Screening for biosurfactant-producing strains

Isolates were screened using two complementary surface-activity assays. In the drop-collapse assay, culture broth was centrifuged at 7500 x g for 20 minutes and 10 µL of cell-free supernatant was deposited on parafilm-coated glass slides and sterile microtiter plates, with distilled water as a negative control; droplet collapse within 30 seconds indicated surface activity (15,16). In the oil-spreading test, 20 µL of cell-free supernatant was applied to a motor-oil film on water, and the diameter of the resulting clear zone was measured; a displacement zone ≥ 1 cm was considered positive (17). Only isolates positive in both assays were retained. These comprised 25 strains, 18 from Qaidam basin (QB01–QB18) and 7 marine (MR01– MR07) (Table 1). Of these, twenty-two were carried into the temperature-only production experiments, and subset of 11 into the combined temperature x salinity experiments.

**Table 1.**
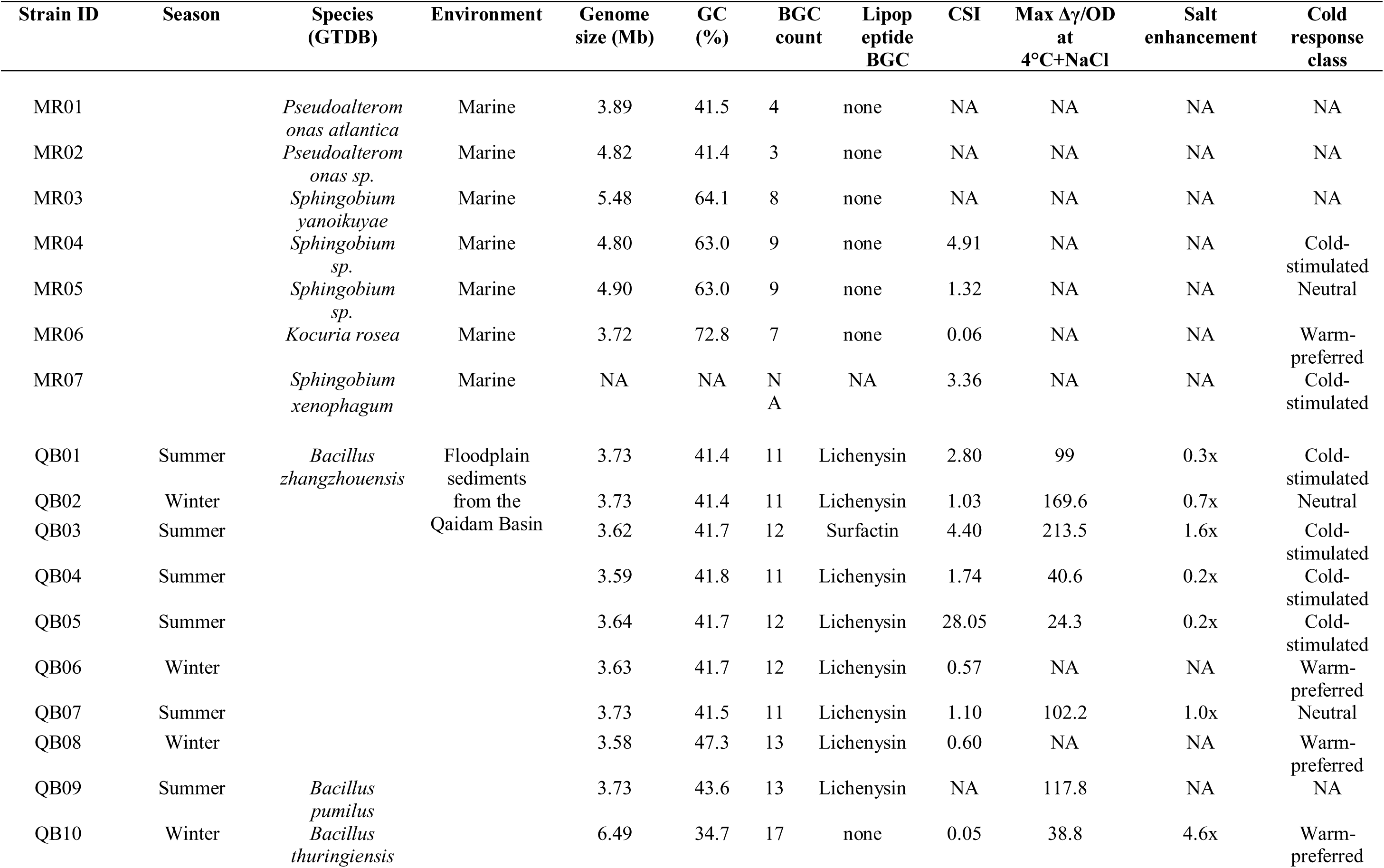
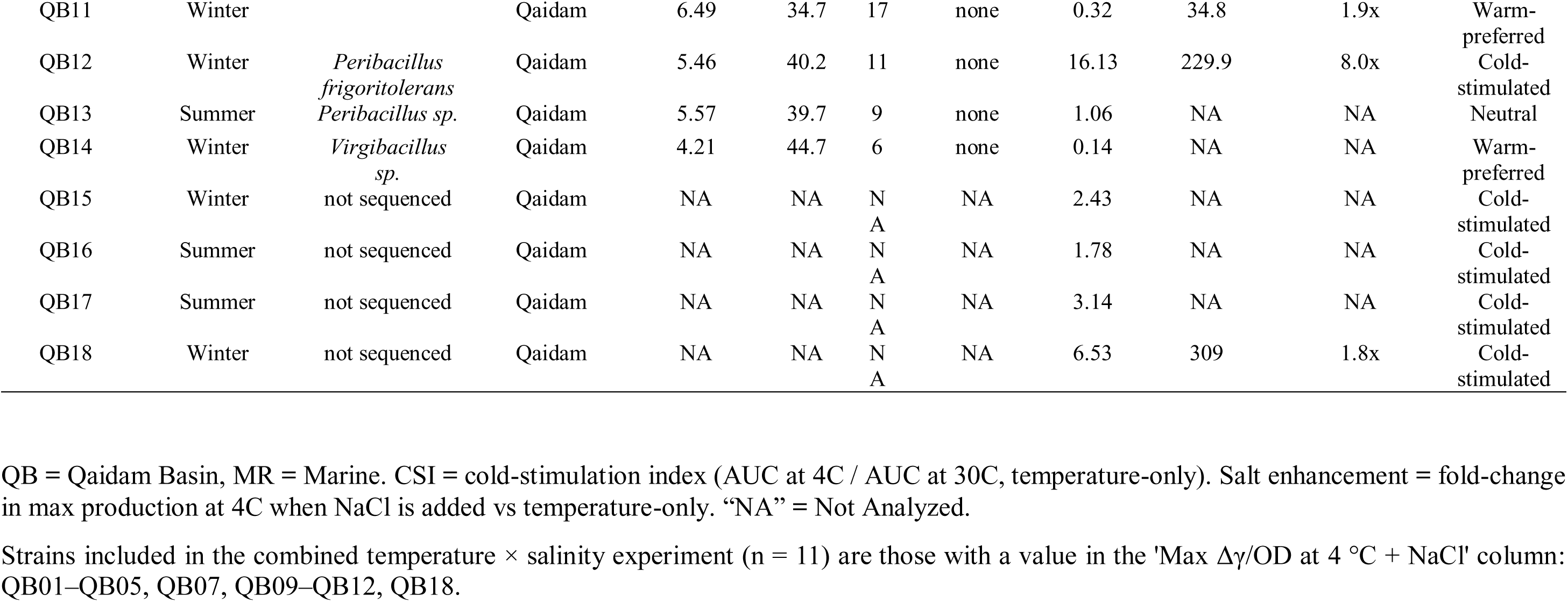
Strain collection, cross-reference identifiers, and key characteristics.

### Biosurfactant simulation experiments

Two complementary experiments were conducted to assess the effects of temperature, salinity, and their interaction on stimulating biosurfactant production.

#### Temperature-only experiments

The 22 biosurfactant-producing strains selected for physiological characterization were cultured in R2A broth at three temperatures (4 °C, 15 °C, and 30 °C), representing cold, sub-optimal, and mesophilic conditions spanning the thermal range of the Qaidam Basin. For each strain × temperature combination, triplicate 250 mL Erlenmeyer flasks containing 150 mL of R2A broth were inoculated from a common LB-agar-revived colony and incubated with shaking at 160 rpm for 11 days.

#### Combined temperature × salinity experiments

A subset of 11 strains (QB01–QB05, QB07, QB09–QB12, and QB18) was further tested across a full factorial design crossing the same three temperatures with three NaCl concentrations (0.5, 1.0, and 1.5 M). For each strain, nine conditions (3 temperatures × 3 salinities, i.e. 9 flasks) were prepared: flasks were distributed among incubators set to the three target temperatures, with all three salinity levels represented at each temperature. R2A broth was supplemented with the appropriate NaCl concentration prior to autoclaving. Inoculation, flask volume (150 mL in 250 mL flasks), shaking speed (160 rpm), and incubation duration (11 days) were identical to the temperature-only experiments.

### Surface tension measurement and biosurfactant quantification

At each of six time points (days 1, 3, 5, 7, 9, and 11), 22 mL of culture was withdrawn from each flask. Optical density at 600 nm (OD₆₀₀) was measured immediately in triplicate and averaged. The remainder was centrifuged at 7,500 × g for 25 min at 4 °C to obtain cell-free supernatant. Surface tension (γ, mN m⁻¹) was measured using a tensiometer (MC-1021 Mince Instrument Equipment Co., Ltd, Xiamen, China), calibrated daily against distilled water (γ = 72.0 ± 0.2 mN m⁻¹ at 25 °C). For each sample, 20 mL of supernatant was analysed in triplicate, and the mean value was used. Uninoculated R2A broth matched to each salinity condition served as the baseline control.

Biosurfactant production was quantified as the reduction in surface tension relative to the uncultured (sterile) control medium:

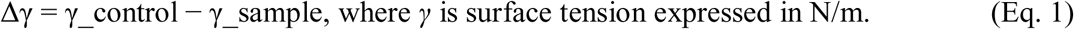

Cumulative biosurfactant activity was calculated as the area under the Δγ-versus-time curve (AUC) using the trapezoidal rule (18) and summed across the five two-day intervals.

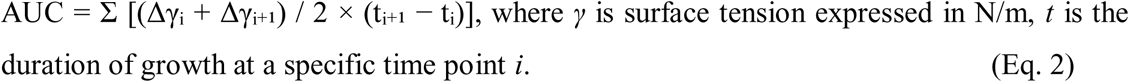

As biosurfactant productivity could increase with increasing biomass, to account for such biomass effect, we introduce specific biosurfactant productivity and define it as Δγ/OD, where Δγ is the reduction in surface tension relative to the control and OD is the optical density measured at 600 nm on the same day. Because Δγ saturates near the critical micelle concentration, this index is a semi-quantitative measure of per-biomass surface activity rather than an absolute yield. Δγ/OD was computed for each individual time point (day-matched) rather than by normalizing all time points to a single whole-experiment value.

### Classification by production capacity

To classify strains by production capacity without imposing arbitrary thresholds, a Gaussian finite mixture model was fitted to the distribution of the per–strain, per–condition maximum Δγ/OD values pooled across all strains and conditions, with the optimal number of components selected by the Bayesian Information Criterion using the mclust package (19). Component boundaries yielded data-driven thresholds for Low, Moderate, and High producer classes (Fig. S1).

### Cold-Stimulation Index

To summarize each strain’s responsiveness to low temperature, we defined a Cold-Stimulation Index (CSI) was as:

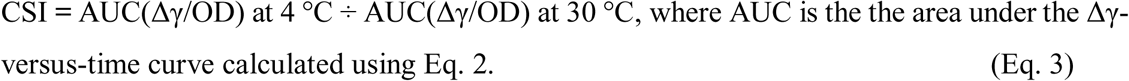

Data obtained from the temperature-only experiments was used to calculate CSI. Strains were categorized as cold-stimulated (CSI > 1.5), neutral (0.7 ≤ CSI ≤ 1.5), or warm-preferred (CSI < 0.7).

### Whole-genome sequencing, assembly, and annotation

Strains selected for genomic analysis were cultured to late exponential phase (7–9 days) in R2A broth for Qaidam and marine isolates, except that MR01, MR02 and MR03 were grown in M2216. Cells were harvested by centrifugation at 7,500 × g for 15 min, washed twice with sterile phosphate-buffered saline solution (PBS, pH 7.4), and the resulting cell pellets were shipped in cold condition to Guangdong Magigene Biotechnology Co., Ltd. (Guangdong, China) for DNA extraction, library preparation, and paired-end whole-genome sequencing.

Raw reads were quality-trimmed with fastp v0.20.1 (20) and assembled *de novo* with SPAdes v3.15.4 in careful mode, which applies mismatch correction and an automatic coverage cut-off to reduce assembly errors (21). These were isolate (pure-culture) genome assemblies, not single-cell or metagenomic assemblies. Contigs shorter than 500 bp were removed. Assembly quality was assessed using QUAST v5.3.0 (22), and genome completeness and contamination were estimated using CheckM2 v1.1.0 (23). In total, 20 high-quality genomes were constructed (100% completeness, ≤ 2.2% contamination; Table 1).

### Taxonomic classification and functional annotation

Genome-based taxonomy was assigned with GTDB-Tk v2.3.0 against GTDB release R214 (24,25). Pairwise average nucleotide identity (ANI) was computed with FastANI v1.34 (26) using the 95% ANI threshold for species delineation. Protein-coding genes were predicted with Prokka v1.15.6 (27) and functionally annotated with eggNOG-mapper v2.1.12 against eggNOG v5.0.2 (28).

### Biosynthetic gene cluster identification

Secondary-metabolite biosynthetic gene clusters (BGCs) were identified with antiSMASH v7.1.0 (2) with KnownClusterBlast, ClusterCompare against MIBiG, and all detection modules enabled. BGC density was calculated as total BGC count per Mb of assembled genome. Gene-cluster families were delineated with BiG-SCAPE v1.1.8 in automatic mode at a raw-distance cut-off of 0.30 (29).

### Pan-genome and phylogenomic analysis

A pan-genome of nine closely related *Bacillus* isolates (*B. zhangzhouensis* QB01-QB08 and *B. pumilus* QB09) was constructed with Panaroo v1.6.0 in moderate cleaning mode (core-genome threshold 0.95) (30). A maximum-likelihood phylogeny was inferred from the core-gene alignment (generated within Panaroo using MAFFT; 3,186 core genes, 2,898,919 aligned nucleotide positions across the nine *Bacillus* isolates, 91.3% invariant sites)with IQ-TREE v2.3.5 (31) under the TIM3+F+I+G4 model selected by ModelFinder (32), with 1,000 ultrafast bootstrap replicates (33).

### Biosurfactant extraction for LC-MS analysis

To obtain lipopeptide-enriched extracts for untargeted metabolomics, strains were cultured in 100 mL of R2A or M2216 broth in 250 mL Erlenmeyer flasks at 15 °C with shaking (160 rpm) for 7 days. This temperature was chosen to balance the adequate biomass for extraction with a sub-optimal temperature relevant to the cold-stress phenotype. After sufficient growth, cultures were acidified to pH 2.0 with 6 M HCl and incubated overnight at 4 °C to precipitate lipopeptides. The acidified cultures were centrifuged at 7,500 × g for 30 min at 4 °C, and the supernatant was carefully decanted. Strains yielding no visible precipitate were classified as extraction-negative and excluded from LC-MS analysis. The resulting pellet was washed twice by resuspension in 10 mL of pH 2.0 distilled water followed by centrifugation at 7,500 × g for 30 min. The washed pellet was dissolved in 2 mL of methanol by thorough vortexing, transferred to a pre-weighed glass vial, and dried under a gentle nitrogen stream. The mass of the acid-precipitated, lipopeptide-enriched crude extract was recorded, and dried extracts were reconstituted in LC-MS-grade methanol at a concentration of 100 µg mL⁻¹ immediately before analysis. This acid-precipitation approach selectively enriches cyclic lipopeptides of the Surfactin/Lichenysin and Fengycin families, which become insoluble below their isoelectric point (34).

### LC-MS/MS analysis

Reconstituted extracts were submitted to LC-MS facility of the Institute of Deep-sea Science and Engineering, Chinese Academy of Sciences in Yazhou Bay, Sanya for untargeted metabolomic profiling. Samples were analyzed on a Thermo Orbitrap system equipped with a heated electrospray ionization source operating in positive-ion mode (H-ESI+). Chromatographic separation, ionization parameters, and MS/MS acquisition settings were determined by the facility. Raw data files (.raw) were converted to open-format mzML using ThermoRawFileParser (35).

### Molecular networking and compound identification

Tandem mass spectra were analysed through the Global Natural Products Social Molecular Networking (GNPS) platform using the METABOLOMICS-SNETS-V2 workflow (36) with a precursor-ion mass tolerance of 2.0 Da, a fragment-ion mass tolerance of 0.5 Da, and default network-filtering parameters (Table S1). Automated dereplication was performed with DEREPLICATOR+ (37), which predicts MS/MS spectra from structural databases and scores spectral similarity to known natural products.

Compound identifications were assigned confidence levels following the approach of Schymanski et al. (2014) (38): Level 1 (confirmed by reference standard), Level 2 (probable structure based on MS² match and library annotation), Level 3 (tentative candidate based on mass match and partial MS² evidence), and Level 4 (unequivocal molecular formula only). For lipopeptide candidates, structural identity was further validated by manual inspection of MS² spectra for diagnostic peptide-backbone fragmentation, specifically, the Leu/Ile sequence ladder (+113.08 Da intervals) characteristic of the Surfactin/Lichenysin heptapeptide (39). Spectral clusters showing non-peptidic fragmentation fingerprints (mass defects < 0.1 Da in the high-mass region, shared fragments across clusters with different precursor masses) were flagged as putative contaminants and excluded after cross-validation against the GNPS contaminant spectral library (Table 2; Fig. S2) (40).

**Table 2.**
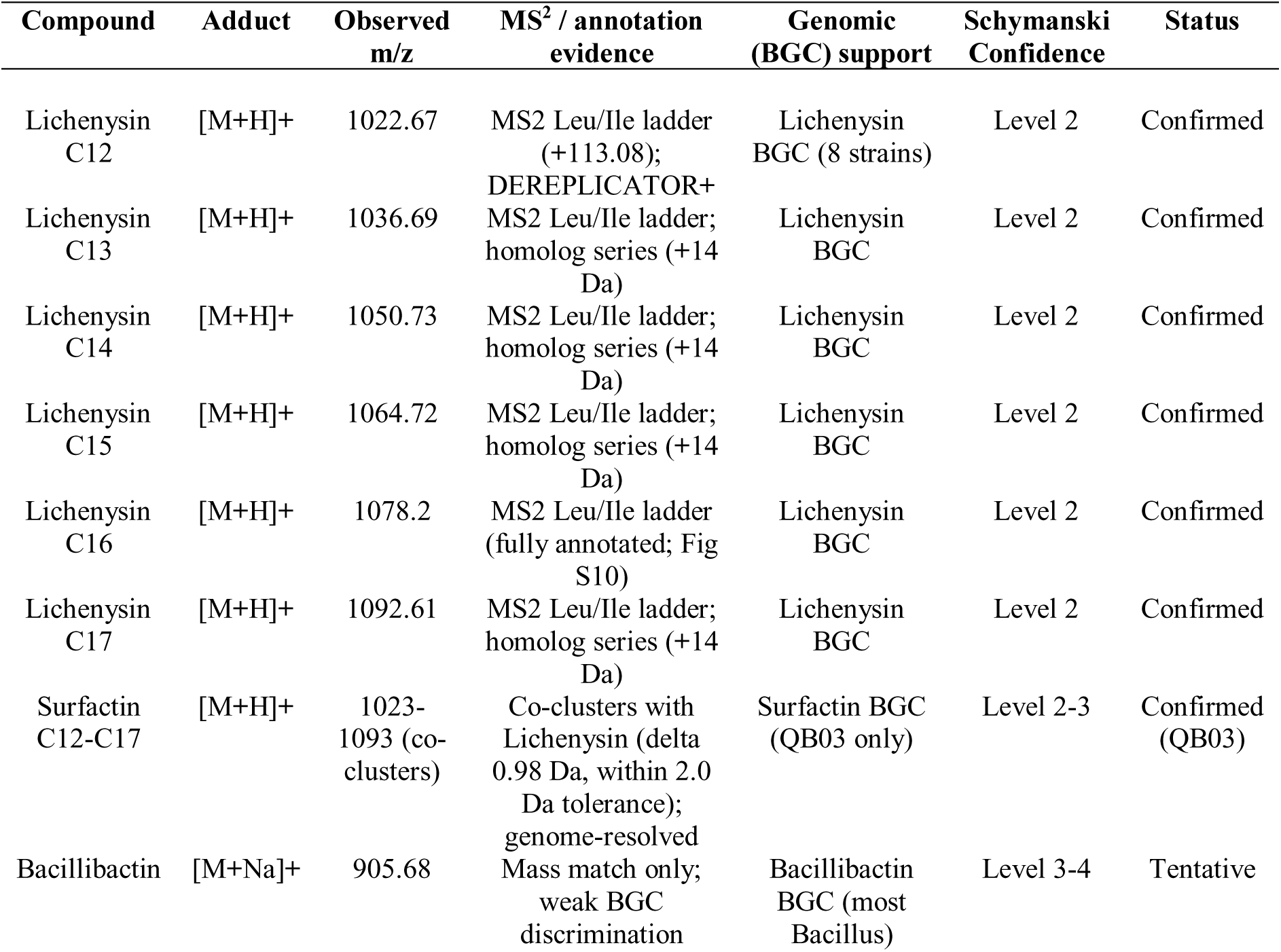
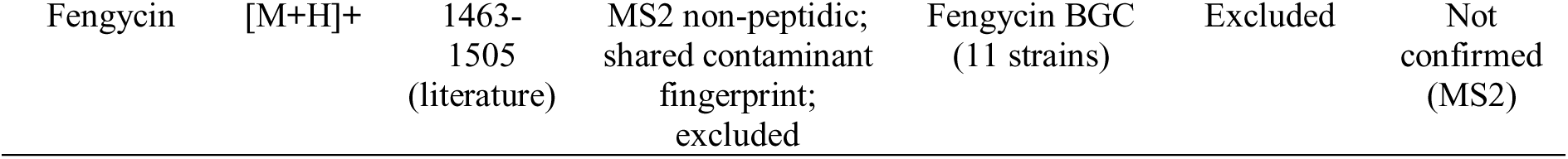
Identified and predicted biosurfactant compounds with Schymanski confidence levels.

Because Lichenysin (containing Gln) and Surfactin (containing Glu) differ by only 0.98 Da and co-cluster at the 2.0 Da precursor tolerance used in this study, these structural isomers could not be distinguished by MS alone. Attribution was therefore resolved genomically: production was assigned as Lichenysin in strains carrying only a Lichenysin BGC and as Surfactin in QB03, the sole carrier of a Surfactin BGC (Table 2).

### Statistical analysis

All statistical analyses were performed in R v4.4.3 (41) with the tidyverse ecosystem for data handling (42), and in Python v3.11 using scipy and scikit-bio for multivariate and phylogenetic analyses.

The effect of temperature on biosurfactant production was tested using a linear mixed-effects model (LMM) fitted by restricted maximum likelihood with lme4 (43), with log₁₀-transformed maximum Δγ/OD as the response, temperature as a fixed effect, and random intercepts for strain and experimental batch. Type III F-tests with Satterthwaite degrees-of-freedom approximation were obtained via lmerTest (44). Marginal and conditional R² were computed following Nakagawa and Schielzeth (2013) (45). Pairwise contrasts with Tukey adjustment and standardized effect sizes (Cohen’s d) were estimated with emmeans (46). For the combined temperature × salinity experiments, salinity and the temperature × salinity interaction were added as fixed effects. A non-parametric Kruskal–Wallis test served as a distributional robustness check.

Genotype–phenotype associations were assessed by Wilcoxon rank-sum tests (with rank-biserial correlation as effect size) comparing cold production between strains with and without lipopeptide BGCs, and by Spearman rank correlations between production and (i) total BGC count and (ii) biosurfactant-regulatory gene count (Table S2). Multivariate differences in COG functional profiles between Qaidam and marine genomes were tested by PERMANOVA (999 permutations) on Euclidean distances using the adonis2 function in vegan v2.7.1 (47,48) (Anderson, 2001). Phylogenetic signal in the cold-response trait was assessed by a Mantel test correlating patristic distances from the core-genome phylogeny with pairwise absolute differences in log₁₀(CSI), with significance evaluated by 9,999 permutations. Growth– production independence was tested by Spearman correlations between maximum OD and maximum Δγ/OD at each temperature (Table S2). Statistical significance was assessed at α = 0.05 throughout, and effect sizes are reported alongside *p*-values to convey biological magnitude independently of sample size (49).

## Results

### Genomic biosurfactant potential is concentrated in the Qaidam *Bacillus* clade

Whole-genome sequencing of 20 isolates (14 Qaidam Basin, 6 marine) yielded high-quality assemblies (100% completeness, ≤2.2% contamination; Table S3) encompassing six genera and eleven species-level groups as classified by GTDB taxonomy (Table 1; Fig. 1A). The Qaidam Basin collection was dominated by a near-clonal clade of eight *Bacillus zhangzhouensis* strains (QB01–QB08; pairwise ANI ≥ 99.40%; Fig. S3), alongside two *Bacillus thuringiensis* (QB10, QB11), one *Bacillus pumilus* (QB09), and representatives of *Peribacillus*, *Virgibacillus*, and a single unsequenced isolate (QB15–QB18). Marine isolates comprised *Pseudoalteromonas*, *Sphingobium*, and *Kocuria* species (MR01–MR06).

**Figure 1.**
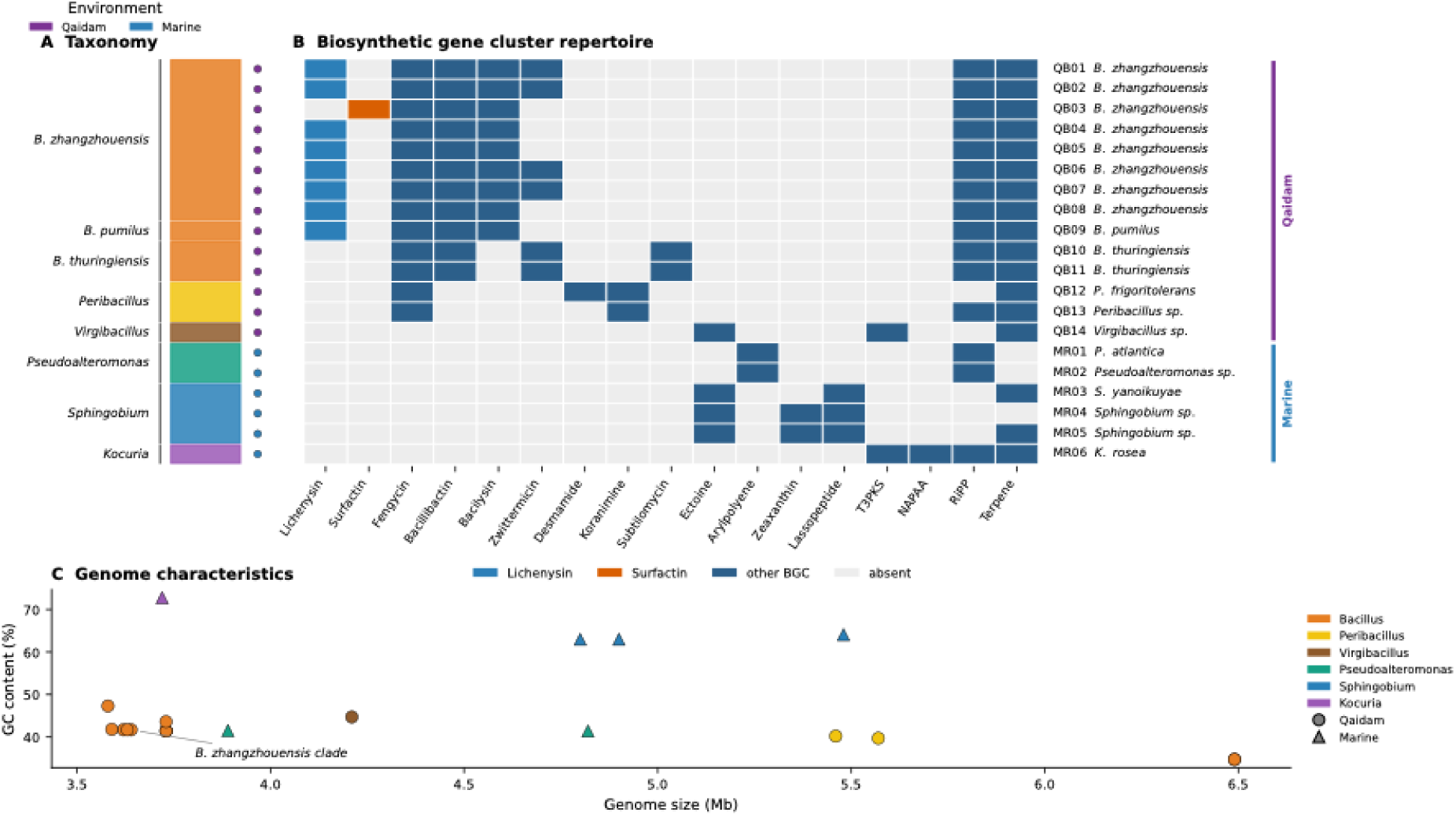
Genomic biosurfactant potential in the 20 studied isolates. (**a**) Taxonomic composition of the 20 sequenced isolates, classified by GTDB and grouped by genus; the colour bar denotes environment of origin (Qaidam Basin, purple; marine, blue). (**b**) Presence/absence of biosynthetic gene clusters (BGCs) across all sequenced genomes (rows, QB01–MR06) for each BGC class (columns), detected by antiSMASH. Lipopeptide clusters are highlighted: Lichenysin (light blue) and the unique Surfactin cluster of QB03 (orange); all other BGCs are shown in dark blue and absent clusters in grey. The right-hand bracket indicates environment (Qaidam, marine). (**c**) Genome characteristics of all isolates, plotting GC content (%) against assembled genome size (Mb); points are coloured by genus and shaped by environment (circles, Qaidam; triangles, marine). The near-clonal *B. zhangzhouensis* clade is indicated.

Genome mining with antiSMASH (2) identified 206 biosynthetic gene clusters (BGCs) across the collection, with a marked enrichment in Qaidam genomes relative to marine genomes (2.80 vs. 1.45 BGCs per Mb; Student’s *t*-test, *p* < 0.01; Table S2). The BGC repertoire differed fundamentally between the two environments: Qaidam *Bacillus* genomes harboured lipopeptide (Lichenysin, Surfactin, Fengycin), siderophore (Bacillibactin), and antibiotic (Bacilysin, Zwittermicin) BGCs, while marine genomes lacked lipopeptide clusters entirely and instead carried Arylpolyene, Ectoine, and Zeaxanthin BGCs characteristic of marine osmotic and oxidative stress adaptations (Fig. 1b; Table S4).

Lipopeptide BGC capacity was tightly concentrated within the *B. zhangzhouensis* clade and *B. pumilus*: all eight *B. zhangzhouensis* strains and QB09 (*B. pumilus*) carried a Lichenysin BGC, while a Surfactin BGC was unique to a single strain, QB03 (Fig. 1b). Fengycin BGCs were predicted in 11 of 14 Qaidam genomes but, as shown below, were not confirmed by LC-MS. Analysis of functional gene content by COG profiling confirmed that environment, rather than individual genome identity, was the dominant structuring factor (PERMANOVA R² = 0.58, *p* = 0.001; Fig. S4; Table S2). Notably, biosurfactant-regulatory genes defined here as genomic copies of the annotated lipopeptide-pathway regulators *degU*, *spo0A*, *phrC* were significantly more abundant in Qaidam genomes than marine genomes (mean 2.93 vs. 0.67 per genome, *p* < 0.001; Table S2), suggesting that the Qaidam environment selects not only for biosurfactant biosynthetic capacity but also with a larger genomic complement of the regulators associated with its expression. Genome characteristics are summarised in Fig. 1c and Table 1.

### LC-MS reveals an extended Lichenysin/Surfactin homolog series as the primary realised biosurfactant

To determine which genomically predicted biosurfactants were actually produced, culture extracts from 14 strains were subjected to untargeted metabolomics by Orbitrap LC-MS/MS and analysed through GNPS molecular networking (36). The analysis yielded 5,726 spectral features grouped into 522 spectral families (Table S1).

Molecular networking identified a dominant spectral family comprising six homologous clusters with precursor masses incrementing by approximately 14 Da ([M+H]⁺: 1022.67, 1036.69, 1050.73, 1064.72, 1078.20, 1092.61; Table 2), consistent with a lipopeptide homolog series differing by successive methylene (CH₂) units in the fatty acid chain (designated C12 through C17). Structural identity was confirmed at Schymanski confidence Level 2 (50) through three independent lines of evidence. First, DEREPLICATOR+ (37) matched the series to Lichenysin/Surfactin-class cyclic lipopeptides. Second, tandem mass spectrometry (MS²) of the C16 homolog (cluster 128273; Fig. 2a) revealed a diagnostic Leu/Ile sequence ladder with four consecutive +113.08 Da intervals (fragment ions at *m/z* 227, 342, 455, 511, 624), confirming the heptapeptide backbone characteristic of the Surfactin family (39). Third, the compound series co-occurred exclusively in strains carrying a Lichenysin or Surfactin BGC (Fig. S5).

**Figure 2.**
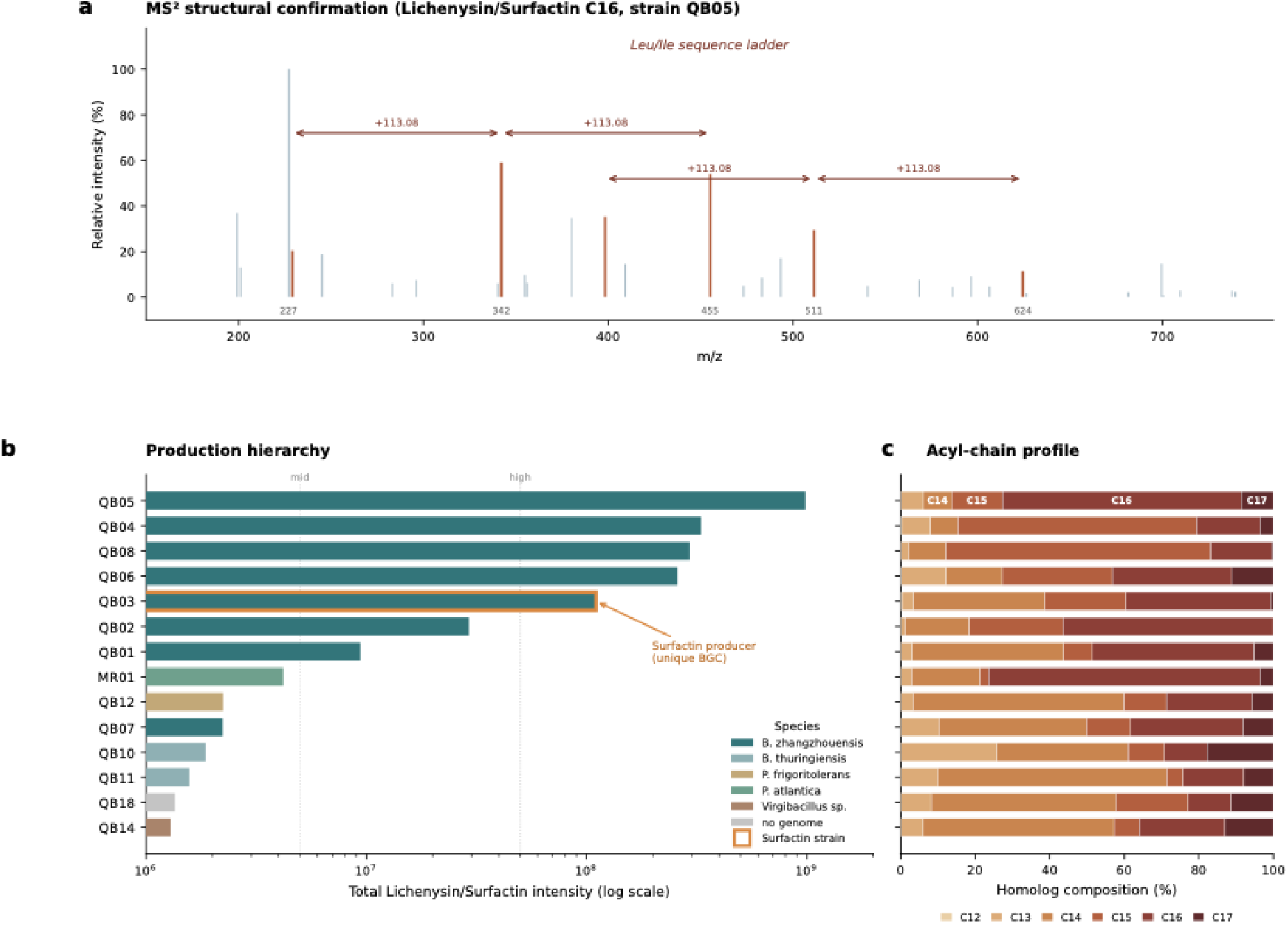
Identity and production of the realised biosurfactants across the strain collection. (**a**) Tandem mass (MS²) spectrum of the C16 homolog (cluster 128273) from strain QB05, as an example, confirming the cyclic lipopeptide backbone. The diagnostic Leu/Ile sequence ladder is annotated, with four consecutive +113.08 Da intervals linking fragment ions at *m/z* 227, 342, 455, 511, and 624. Duplicate LC-MS runs for QB04 and QB08 were averaged to a single value per strain. (**b**) Production hierarchy of the Lichenysin/Surfactin homolog series, shown as a semi-quantitative measure indicating the abundance (log₁₀ scale) summed across the C12–C17 homologs for each LC-MS-analysed strain; bars are coloured by species, and the sole Surfactin-BGC carrier (QB03) is outlined in orange. Dashed vertical lines mark the data-driven mid- and high-production thresholds. (**c**) Acyl-chain composition of the homolog series for each producing strain, shown as the percentage contribution of each homolog (C12–C17) to total Lichenysin/Surfactin intensity (100% stacked bars; warm-to-dark colour ramping from C12 to C17).

Lichenysin (containing Gln) and Surfactin (containing Glu) differ by only 0.98 Da, thus these structural isomers co-clustered at the 2.0 Da GNPS precursor mass tolerance and would obscure their identification. We resolved this ambiguity using genomic data. Seven of the eight *B. zhangzhouensis* strains carried only the Lichenysin BGC and no sign of Surfactin BGC and were therefore assigned as Lichenysin producers, whereas QB03, the sole surfactin-BGC carrier, was assigned as surfactin producer (Table 2). This genome-guided disambiguation illustrates the complementary power of integrating metabolomics with whole-genome sequencing (51).

Production of the Lichenysin/Surfactin series varied by approximately three orders of magnitude across strains (Fig. 2c). The five highest producers, QB05 > QB04 > QB08 > QB06 > QB03, were all *B. zhangzhouensis*, while non-*Bacillus* strains and the two *B. thuringiensis* (QB10, QB11) showed only trace-level of lipopeptide signals. Among the high producers, acyl-chain composition shifted systematically: the top producer QB05 was enriched in longer-chain homologs (C16, C17), whereas lower producers showed a proportionally higher fraction of shorter chains (C14, C15; Fig. 2b), suggesting that higher biosynthetic flux favours chain elongation (52).

Fengycin-range spectral features were genomically predicted in 11 Qaidam strains (Fengycin BGC present), but MS² analysis of the candidate clusters (112611, 112920) revealed non-peptidic fragmentation fingerprints inconsistent with a lipopeptide backbone (Fig. S2). Specifically, the five most intense high-mass fragments shared mass defects below 0.1 Da (e.g., *m/z* 359.029, 415.037, 285.010), far below the 0.2–0.4 Da range expected for peptide bond fragments, and 50–70% of fragment ions were shared across clusters with different precursor masses, a pattern diagnostic of chemical contaminants rather than biologically distinct metabolites (40). These clusters were therefore excluded from further analysis (Table 2). A tentative Bacillibactin annotation ([M+Na]⁺ = 905.68; Level 3–4) was retained based on mass match and genomic support (Table 2; Fig. S5). The compound identification strategy and confidence assessments for all detected biosurfactants are summarised in Table 2.

### Cold and salt stress jointly and strain-specifically shape biosurfactant production

Having established that Lichenysin/Surfactin is the primary realised biosurfactant and that its production is concentrated in the *B. zhangzhouensis* clade, we next tested the central hypothesis: whether extreme environmental conditions, specifically low temperature and high salinity, actively induce biosurfactant production. Twenty-two strains were phenotyped for surface-tension-reducing activity (Δγ/OD) across three temperatures (4, 15, 30 °C) in temperature-only experiments and, for a subset of 11 of these strains, across a full factorial temperature × salinity design (0.5, 1.0, 1.5 M NaCl) (Tables S4, S5).

At the community level, temperature did not significantly affect mean specific biosurfactant productivity (Δγ/OD) (LMM: *F*(2, 45.9) = 1.84, *p* = 0.17; Table S2), and the interaction between temperature and salinity was likewise non-significant (*F*(4, 70.3) = 1.86, *p* = 0.13). However, strain identity accounted for the majority of variance (conditional *R*² = 0.68 vs. marginal *R*² = 0.018), indicating that response to low temperature is not a universal microbial trait but rather a strain-specific capacity. This observation motivated a strain-level analysis.

Individual strain trajectories revealed that of the 21 strains with a defined CSI (QB09 was tested only in the salinity design and has no CSI), 11 were cold-stimulated CSI > 1.5) 6 were warm-preferred (CSI < 0.7) and 4 were neutral; Fig. S6, Table 1; Table S5).

The combined-stress experiment revealed strong, strain-specific effects of added salt on cold production Fig. 3b; Table S2). In several strains, adding salt markedly increased production at 4 °C including QB12 (8.0-fold; 229.9 vs 28.7 Δγ/OD), and also QB10 (4.6-fold), QB11 (1.9-fold), QB18 (1.8-fold), and QB03 (1.6-fold), whereas in others salt suppressed the cold response (e.g. QB01, QB04, and QB05, each ≤ 0.3-fold). The combined-stress data further revealed strain-specific salt-enhancement patterns (Table 1; Table S6; Fig. S8). Notably, QB12 (*Peribacillus frigoritolerans*) showed an 8.0-fold increase in surface-tension-reducing activity at 4 °C + 1.0 M NaCl relative to 4 °C alone (229.9 vs. 28.7 Δγ/OD), making it the second most active strain under combined stress despite lacking any lipopeptide BGC (implying a non-lipopeptide surfactant; Table 1). Several strains including QB04 and QB05 achieved peak production not at 4 °C but at 15 °C + 1.5 M NaCl (263.9 and 278.3 Δγ/OD, respectively; Table S6), indicating that the optimal stress condition for maximal biosurfactant output is strain-dependent and does not always correspond to the most extreme cold.

**Figure 3.**
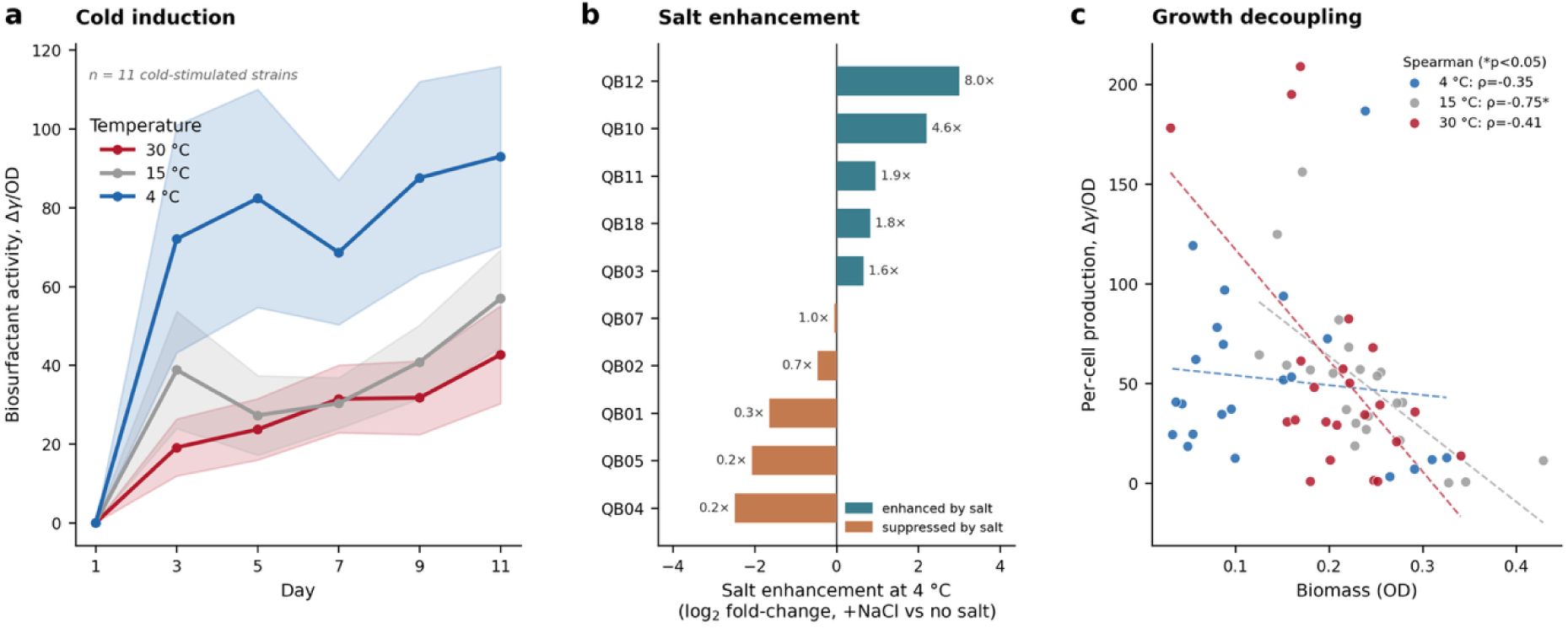
Cold and salt stress synergistically and strain-specifically induce biosurfactant production. (**a**) Time course of specific biosurfactant activity (Δγ/OD) over the 11-day incubation for the cold-stimulated strains, shown as mean ± standard error at each of three temperatures (4 °C, blue; 15 °C, grey; 30 °C, red). (**b**) Salt enhancement of the cold response, shown for the strains tested under combined stress as the fold-change in maximum specific productivity (Δγ/OD) at 4 °C with added NaCl relative to 4 °C without added salt; values above 1 indicate that added salt increased cold production (e.g. QB12, 8.0-fold), values below 1 indicate suppression (**c**) Relationship between specific biosurfactant productivity (Δγ/OD) and biomass (OD) at each temperature, with Spearman correlations (4 °C, ρ = −0.35; 15 °C, ρ = −0.75, *p* < 0.001; 30 °C, ρ = −0.41); the negative associations indicate that production is independent from growth. Points are coloured by temperature as in (a).

A hallmark of an actively induced stress responses, as opposed to passive metabolic overflow, is that production is independent from growth. At all three temperatures, per-cell biosurfactant production was negatively correlated with biomass (Spearman *ρ* = −0.35, −0.75, −0.41 at 4, 15, 30 °C, respectively; significant at 15 °C, *p* < 0.001, and 30 °C, *p* = 0.05; Fig. 3c; Fig. S9; Table S2). At 4 °C, growth was most suppressed yet per-cell production was highest, directly contradicting a biomass-driven model and supporting active induction under cold stress.

### Regulatory-gene content, not biosynthetic-cluster number, predicts biosurfactant production

To link genomic features to the observed phenotypic variation, we integrated whole-genome data with cold-production metrics across the 17 strains for which both genome and physiology data were available (Table 1).

The strongest genomic predictor of cold biosurfactant production was the genomic copy number of biosurfactant-regulatory genes (*degU*, *spo0A*, *phrC*), a genome content measure (Spearman *ρ* = 0.85, *p* < 0.001, *n* = 17; Fig. 4a; Table S2). This relationship was stepped rather than linear: strains with 0–1 regulatory genes (exclusively marine) showed little specific productivity at 4 °C (median ∼35 Δγ/OD), those with 2 produced moderately (∼39), and those with 3–4 (exclusively Qaidam *Bacillus*) produced substantially more (median ∼194; Fig. 4a). Because regulatory-gene count covaried with ecological origin, this finding reflects both a genuine gene-dosage effect and the broader Qaidam enrichment in biosurfactant capacity (see Discussion).

**Figure 4.**
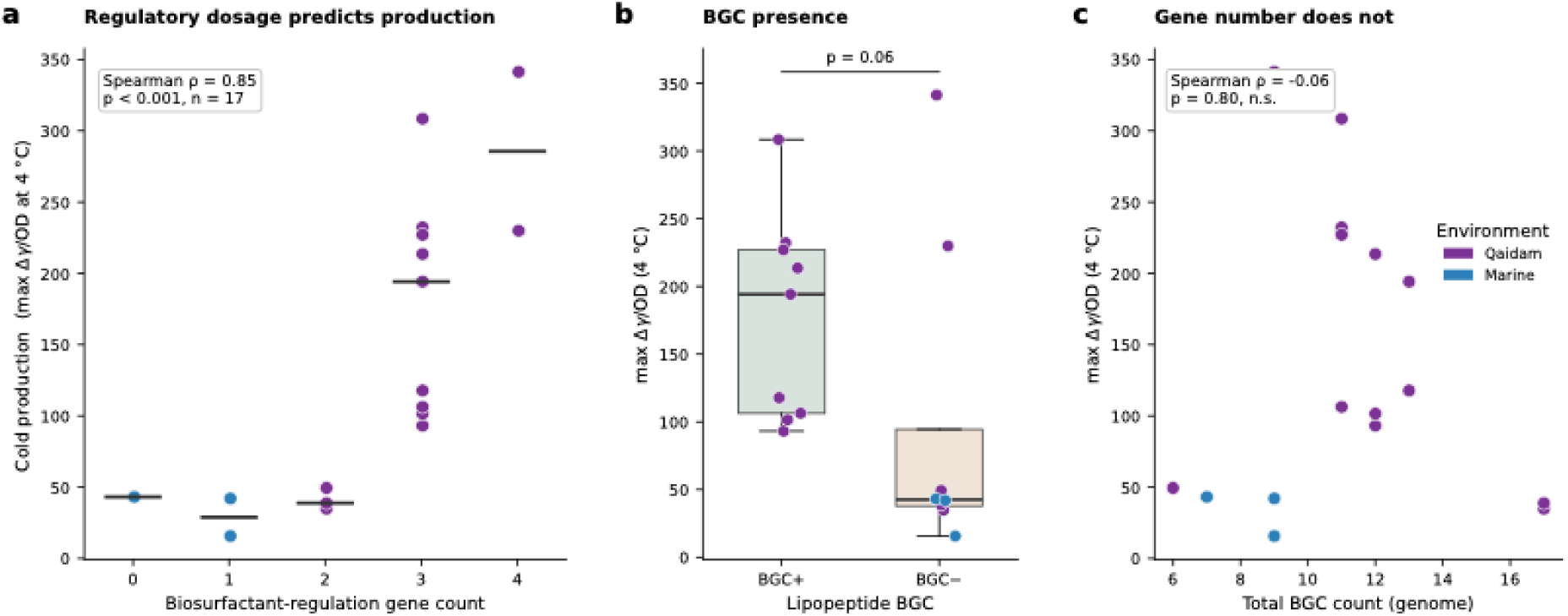
Regulatory capacity, not biosynthetic-cluster number, predict biosurfactant production. (**a**) Cold biosurfactant production (maximum Δγ/OD at 4 °C) as a function of the number of biosurfactant-regulatory genes per genome (*degU*, *spo0A*, *phrC*); the relationship is strong and positive (Spearman ρ = 0.85, *p* < 0.001, *n* = 17). Horizontal bars denote group medians; points are coloured by environment (Qaidam, purple; marine, blue). (**b**) Cold production compared between strains carrying a lipopeptide BGC (BGC+) and those lacking one (BGC−); the two BGC-strains that mount a cold response, QB12 (*Peribacillus frigoritolerans* and MR04 (*Spingobium sp.))* box plots show median and interquartile range with individual strains overlaid (Wilcoxon rank-sum *W* = 16, *p* = 0.06, rank-biserial *r* = 0.56). (**c**) Cold production as a function of total BGC count per genome, showing no relationship (Spearman ρ = −0.06, *p* = 0.80, *n* = 17). All the three panels show that genomic content of regulatory genes, but not total biosynthetic-cluster count, predicts realised production.

Lipopeptide-BGC presence was associated with higher cold production (median 194 vs. 43 Δγ/OD for BGC+ vs. BGC−; Wilcoxon *W* = 16, *p* = 0.06, rank-biserial *r* = 0.56; Fig. 4b), and Qaidam strains produced significantly more than marine strains at 4 °C (median 156 vs. 42 Δγ/OD; Mann–Whitney *U* = 38, *p* = 0.03; Table S2).

Critically, total BGC count (all classes) per genome, the crudest measure of biosynthetic potential, showed no relationship with production (*ρ* = −0.06, *p* = 0.80; Fig. 4c; Table S2). Strains with 17 BGCs (QB10, QB11) produced less than strains with 11 (QB01, QB05). This null result, juxtaposed with the strong regulatory-gene association, demonstrates that it is the specific regulatory network controlling biosurfactant expression, rather than the sheer genomic capacity of biosynthetic pathways, that determines the functional cold-stress phenotype among the strains examined.

### The cold-stress biosurfactant response is strain-specific, and biochemically heterogeneous

The preceding sections established that genomic potential (Section 3.1), realised chemistry (Section 3.2), and stress-induced phenotype (Section 3.3) all point to the *B. zhangzhouensis* clade as the primary biosurfactant-producing lineage. To test whether these three levels of evidence converge at the individual-strain level, and whether the cold response is a phylogenetically conserved clade trait or a strain-specific regulatory phenomenon, we aligned genotype, chemotype, and phenotype data on the core-genome phylogeny of the *B. zhangzhouensis* clade (Fig. 5a).

**Figure 5.**
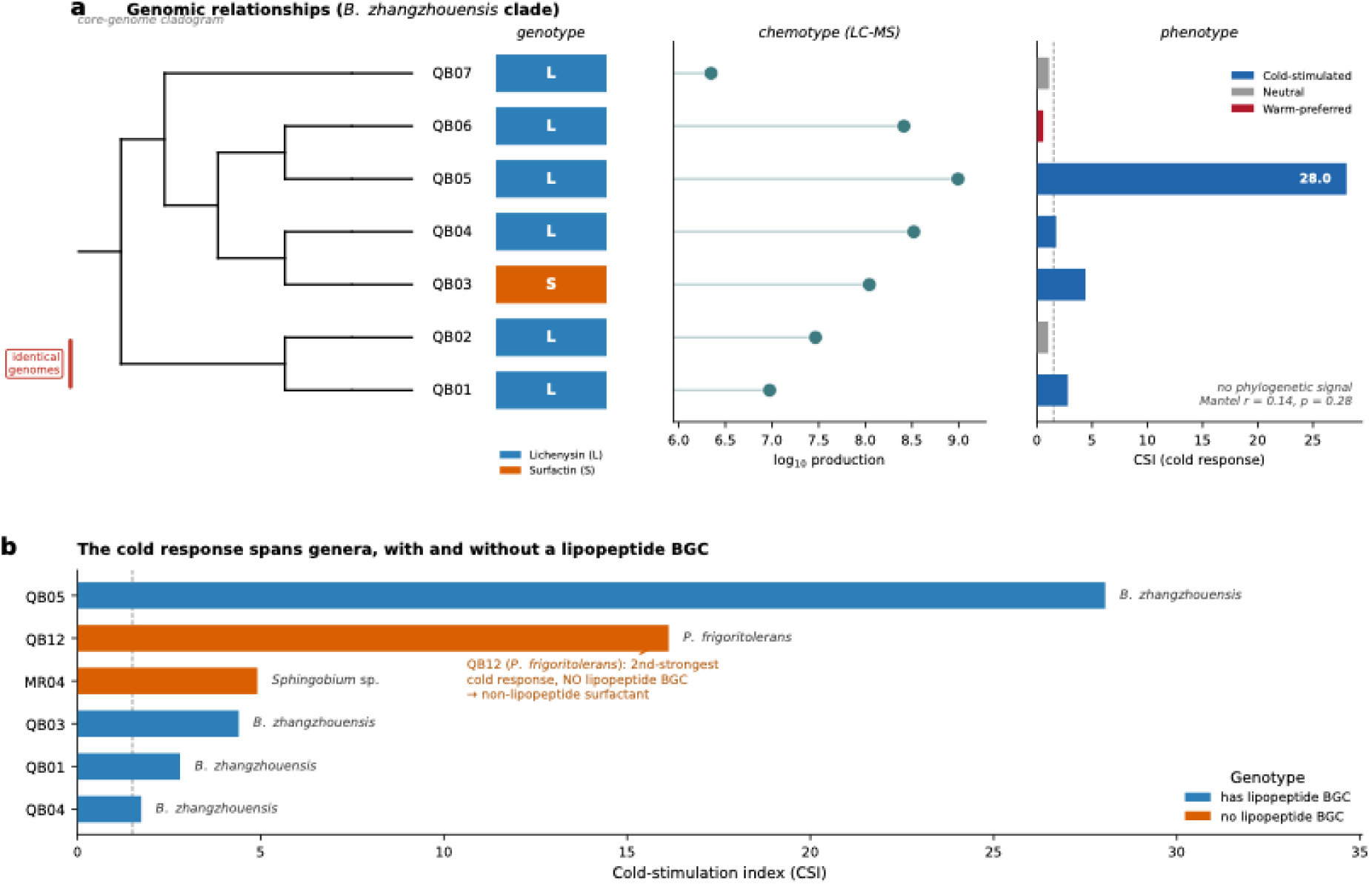
Biosurfactant production is a regulated, strain-specific response to cold stress, independent from phylogeny. (**a**) Integration of genotype, chemotype, and phenotype across the near-clonal *B. zhangzhouensis* clade, aligned on the core-genome maximum-likelihood cladogram (left). The *genotype* track shows the lipopeptide BGC carried by each strain (Lichenysin, blue, “L”; Surfactin, orange, “S”); the *chemotype* track shows realised lipopeptide production (log₁₀ LC-MS intensity, lollipops); and the *phenotype* track shows the cold-stimulation index (CSI), coloured by cold-response class (cold-stimulated, blue; neutral, grey; warm-preferred, red). The clonal pair QB01/QB02 (effectively identical genomes) is bracketed; the cold-response trait shows no phylogenetic signal (Mantel *r* = 0.14, *p* = 0.28). (**b**) The cold-stress response spans three genera and occurs both with and without a lipopeptide BGC; CSI is shown for the genome-sequenced cold-stimulated strains, coloured by lipopeptide-BGC presence (blue, present; orange, absent). QB12 (*Peribacillus frigoritolerans*), the second-strongest cold responder, carries no lipopeptide BGC, implying a non-lipopeptide surfactant. Dashed line, cold-stimulation threshold (CSI = 1.5).

Within this near-clonal clade (all pairwise ANI > 99.4%; Fig. S3), the lipopeptide BGC genotype was highly conserved: six of seven strains carried Lichenysin, one (QB03) carried the unique Surfactin BGC (Fig. 5a). Chemotype (LC-MS lipopeptide production) varied over approximately three orders of magnitude but was universally above background in all clade members. In contrast, the cold-response phenotype (CSI) was not phylogenetically structured.

CSI ranged from 0.57 (QB06, warm-preferred) to 28.05 (QB05, the strongest cold-responder in the collection), and a Mantel test confirmed no significant correlation between patristic distance and cold-response divergence (Mantel *r* = 0.14, *p* = 0.28, *n* = 7; Fig. 5a). Two comparisons make the point directly. First, the clonal pair QB01 and QB02 (patristic distance 6.5 × 10⁻⁸ substitutions per site) fell on opposite sides of the cold-stimulation threshold (CSI = 2.80, cold-stimulated, vs. 1.03, neutral; Fig. 5a). Second, QB05 and QB06 — both *B. zhangzhouensis* carrying the same lichenysin BGC — differ ∼49-fold in CSI (28.05 vs. 0.57), spanning the full range from strongest cold-responder to warm-preferred. These results demonstrate that cold-inducible biosurfactant production within the clade is a regulatory-level trait, not a lineage-level trait: strains with near-identical genomes and identical biosynthetic gene clusters can mount fundamentally different stress responses.

Beyond the *B. zhangzhouensis* clade, the cold-stress biosurfactant response was chemically heterogeneous (Fig. 5b). Among the six genome-sequenced cold-stimulated strains, four carried a lipopeptide BGC (*B. zhangzhouensis*) while two, QB12 (*Peribacillus frigoritolerans*, CSI = 16.13) and MR04 (*Sphingobium* sp., CSI = 4.91), lacked any lipopeptide BGC entirely (Fig. 5b; Table 1). QB12’s case is particularly interesting: it was the second-strongest cold-responder across the entire collection and showed an 8.0-fold salt-enhancement at 4 °C, yet its LC-MS lipopeptide signal was ∼440-fold lower than the top lipopeptide producer QB05 (Table 1). The surface-tension assay employed in the physiology experiments is class-agnostic, it detects any molecule that reduces surface tension, regardless of chemical class, whereas the ESI-positive-mode LC-MS workflow preferentially ionises lipopeptides. The discordance between strong surface-tension activity and negligible lipopeptide signal in QB12 therefore indicates the production of a non-lipopeptide surfactant (possibly a glycolipid or phospholipid derivative) under cold-salt stress, a finding that would have been invisible without the multi-method design (53). The converse pattern, a strong LC-MS lipopeptide signal in a strain lacking a genomically detected lipopeptide BGC, was not observed, making it unlikely that QB12’s activity reflects a lipopeptide BGC missed by antiSMASH.

## Discussion

Biosurfactant production in the Qaidam Basin isolate collection is a regulated, strain-specific response to cold-saline stress rather than a fixed property of any single lineage, and the capacity to mount this response is distributed across the community in ways that genome studies alone cannot predict. By integrating comparative genomics, untargeted metabolomics, and quantitative physiology across the same isolate set, we resolved biosurfactant production into three separable tiers of control: genomic capacity, which sets the outer boundary of what a strain can synthesise; regulatory approach, which determines how readily that capacity is mobilised under stress; and strain-level regulatory state, which dictates what is actually produced. The central finding, that these tiers are controlled by different determinants, and that the realised phenotype is independent from both phylogeny and biosynthetic-gene composition, redefine a question that has long constrained the study of microbial natural products in extreme environments: not whether an organism *can* produce a stress-responsive surfactant, but what controls whether it *does*. On our data the answer has two parts: at the leavel of gene content, the presence of the regulator complement (*degU*, *spo0A*, *phrC*), not the size of the BGC repertoire, markers which strains are capable of a strong response; and among strains sharing that complement, the amount produced is set downstream of gene content, at each strain’s physiological state (the clonal-pair results show it directly). This distinction has direct consequences for how biosurfactant-producing taxa, and by extend, production of other BGC metabolites, should be prospected and predicted, in cold, hypersaline habitats and other stresses.

The most immediate implication concerns the interpretation of genomic potential. We found that the number of biosurfactant-regulatory genes per genome strongly predicted cold biosurfactant production (Spearman ρ = 0.85, p < 0.001), whereas the total number of biosynthetic gene clusters, the most commonly invoked proxy for biosynthetic potential in genome-mining studies, showed no relationship (ρ = −0.06, p = 0.80). This dissociation is consequential because BGC counts are routinely used to rank strains and taxa by their presumed metabolic richness, an approach that implicitly assumes biosynthetic capacity translates into output (REF?). Our data show that, at least for stress-responsive biosurfactants, this assumption fails: strains carrying the largest BGC repertoires (the *B. thuringiensis* isolates, with 17 clusters each) were among the weakest producers, while the genomic feature that best predicted production was the regulator-gene complement. This finding aligns with the broader recognition that a substantial fraction of microbial BGCs are transcriptionally silent under standard conditions (53) and suggest that the genotype-to-phenotype gap in secondary metabolism as, in part, a question of regulatory capacity rather than biosynthetic repertoire. We do not claim this as the whole explanation, culture conditions, promoter and regulatory-sequence variation, and post-transcriptional control also determine whether a BGC is expressed, but for the targeted discovery of cold-active biosurfactant producers, our results argue that regulatory-gene content is a more informative genomic marker than BGC abundance.

If regulatory capacity controls production, then phenotype should be separable from phylogeny even among closely related strains, and this is precisely what we observed. Within a near-clonal *Bacillus zhangzhouensis* clade sharing greater than 99.4% average nucleotide identity, the cold-stimulation phenotype showed no phylogenetic signal (Mantel r = 0.14, p = 0.28), and the clonal pair QB01 and QB02, separated by a patristic distance of only 6.5 × 10⁻⁸ substitutions per site and carrying identical lichenysin biosynthetic and regulatory gene complements, fell on opposite sides of the cold-stimulation threshold (CSI = 2.80 versus 1.03). Because these strains possess the same biosynthetic operon and the same regulatory genes, the divergence in their realised phenotype cannot be explained at the level of gene regulatory activity or physiological state. We did not measure transcription or regulatory activity; this mechanism is an inference consistent with our data and with well-characterised control of lipopeptide biosynthesis in *Bacillus*. In that model system, production is gated not by the synthetase operon itself but by a signal-integrating regulatory network: the quorum-sensing response regulator ComA activates the *srfA*/lichenysin operon in response to the ComX pheromone (54,55), while DegU and the sporulation master regulator Spo0A further modulate output in response to nutrient and stress cues, with DegU acting as a direct repressor of the lichenysin *lchA* operon in *B. licheniformis* (56,57). Because the realised level of lipopeptide synthesis depends on the phosphorylation state and integrated activity of this network rather than on the mere presence of its components, two strains with identical regulatory genes can produce markedly different amounts under the same conditions, through differences in signal thresholds, stochastic variation in regulator expression, or sub-ANI sequence variation in regulatory or promoter regions invisible to genome-level comparison. Even as an inference, the clonal-pair divergence is therefore not an anomaly but an expected outcome if the trait is controlled at regulatory rather than structural-genetic level, and it shows how accurately genome sequence alone can predict stress-responsive secondary-metabolite phenotypes (58).

The stress-responsive surfactant phenotype was, moreover, biochemically heterogeneous and convergent across genera rather than confined to a single biosynthetic class. While lipopeptide production dominated the *B. zhangzhouensis* clade, the second-strongest cold responder in the entire collection, QB12 (*Peribacillus frigoritolerans*, CSI = 16.13) showed an 8.0-fold enhancement of surface activity at 4 °C under salt stress (Table 1), yet its lipopeptide LC-MS signal was 440-fold lower than the top lipopeptide producer. Because the LC-MS workflow is biased toward detecting lipopeptides, a lipopeptide responsible for this activity would have been detected. Its near absence, rather than the absence of a predicted BGC, therefore indicate that surfactant non-lipopeptide in nature. We note that antiSMASH can miss fragmented or divergent clusters, so we do not relay on genome annotation alone to exclude lipopeptide production. The apparent contradiction is resolved by the orthogonal detection profile of two assays: the surface-tension assay is functionally generic, responding to any interfacial-active molecule capable of lowering the surface tension, whereas positive-mode electrospray LC-MS preferentially ionises lipopeptides. The deviation in QB12 strong surface activity with a negative lipopeptide ion signal is therefore not a contradiction but a positive indication that it produces a non-lipopeptide surfactant, possibly a glycolipid or related amphiphile, under combined cold-salt stress. This finding has two broader implications. Biologically, it demonstrates that the cold-stress biosurfactant response is a convergent strategy reached through biochemically distinct routes in different lineages, broadening the cold-stress biosurfactant response beyond the lipopeptide-producing *Bacillus* that dominate the biosurfactant literature. Methodologically, a genome only survey would have assigned QB12 as a non-producer category, and LC-MS alone would have failed to detect any surfactant phenotype. The non-lipopeptide surfactant was identified only through the orthogonal integration of a functional activity assay and targeted metabolite detection (59).

The co-stimulation of biosurfactant production by low temperature and high salinity, with added salt amplifying the cold response up to 8-fold in QB12 reflects a stress-mitigation strategy shaped by the native habitat of these isolates. In the Qaidam Basin, cold and hypersalinity are not independent selective pressures but co-occurring features of the same environment, making a joint response ecologically consistent. The adaptive functions that biosurfactants could possibly serve under such conditions are several and not mutually exclusive: lipopeptides and other amphiphiles can modulate membrane fluidity and integrity under cold and osmotic stress, contribute to cryoprotection and desiccation resistance, enhance access to hydrophobic substrates and nutrients in oligotrophic sediments, and influence biofilm formation and surface motility (60). Lichenysin in particular is notable in this context: relative to surfactin, it exhibits higher surface activity and greater stability under extremes of salinity, pH, and temperature, and it is a more efficient cation chelator (61), properties that would be advantageous in a fluctuating hypersaline environment and that may help explain why the lichenysin pathway, rather than surfactin, predominates in this cold-saline-adapted clade. We emphasize, however, that our data establish that production is stress-responsive and regulated, not that it confers a measured fitness benefit; the adaptive interpretation, though well-motivated, remains an inference pending direct fitness assays.

These observations acquire a broader significance from the provenance of the strains. The Qaidam Basin is one of Earth’s most studied terrestrial analogues for ancient Martian lacustrine environments, combining hyperaridity, intense radiation, evaporite mineralogy, freeze–thaw cycling, and fluctuating hypersalinity in a single environment (62,63). The demonstration that bacteria from such an environment mount a regulated, stress-inducible surfactant response, and that this capacity is distributed across several genera yet expressed in a strain specific manner, but phenotypically individualized, shows how microbial survival strategies might operate under comparable extraterrestrial stresses, and identifies surface-active lipopeptides as candidate molecules whose production is upregulated precisely under the cold, saline conditions thought to characterize habitable niches on early Mars (64). Because lipopeptides such as lichenysin are chemically robust, remaining surface-active across wide range of temperature and salinity (65) and structurally distinctive, their potential as preservable biosignatures in evaporitic sediments warrants consideration (66,67). We propose this connection cautiously and as a motivation for further work rather than a claim, but it situates the regulated biosurfactant response we describe within the larger question of how, and under what controls, life that persists in environments at the edge of habitability.

Several limitations bound these interpretations and define the most productive next experiments. First, our proposed mechanism of the strain-specific and clonal-pair divergence to differential regulatory-network activity, while consistent with the established control of lipopeptide biosynthesis in *Bacillus*, rests on genomic gene content rather than expression data; targeted transcriptomics or reporter assays across temperatures and salinities are needed to confirm that the ComA/DegU/Spo0A network mediates the responses we observe, and provide further details in the regulatory systems. Second, the strong correlation between regulatory-gene content and production covaries with ecological origin, since the regulator-rich genomes are also the Qaidam *Bacillus*; separating gene-dosage effects from broader clade- and environment-level differences will require a more taxonomically balanced sampling. Third, the phylogenetic-signal analysis was necessarily restricted to the seven genome-sequenced members of the *B. zhangzhouensis* clade, and although the clonal-pair result is unambiguous, a larger set of closely related strains would strengthen the generality of the phylogeny– phenotype independence. Finally, the non-lipopeptide surfactant inferred in QB12 remains structurally unidentified; its characterization, through feature-based molecular networking and complementary ionisation modes, is a clear priority and may reveal a previously undescribed class of cold-active surfactant.

Overall, our results support a model in which biosurfactant production in extreme cold-saline environments is determined hierarchically, genomic capacity sets the boundary, regulatory approach sets the responsiveness, and strain-level regulatory state sets the realised output, and is achieved through chemically diverse molecules across multiple genera. This model makes testable predictions: that strains with equivalent biosynthetic genes but differing regulatory-network configurations will diverge predictably in stress-induced production; that the cold-saline surfactant response is transcriptionally rather than genomically encoded and should be reversible and condition-dependent; and that functionally analogous, chemically distinct surfactants await discovery in the non-lipopeptide producers of these communities. More broadly, by showing that the capacity for biochemical adaptation to extreme conditions is distributed across a community in a manner independent from phylogeny and unpredictable from biosynthetic-gene capacity, this work argues that understanding microbial function in extreme environments, on Earth and beyond, requires moving past genomic potential to the regulatory logic that controls whether that potential is ever realised.

## Author Contributions

***Habib Ullah:*** *Conceptualization, data curation, formal analysis, methodology, software, visualization, and writing original draft. Conducted all bioinformatic analyses*.

***Zhuang Han****: Assists in the metabolomic part of the study*.

***Maggie C. Y. Lau Vetter***: Conceptualization, data interpretation, supervision, and project administration. Provided critical intellectual guidance throughout all stages of the study and reviewed the manuscript.

## Deceleration of Competing Interest

The author declares that there is no conflict of interest.

## Acknowledgments

This work was supported by ANSO (Alliance of International Science Organizations) Scholarship for young talent to H.U., and Discretionary Grant (Institute of Deep-sea Science and Engineering; grant number IDSSE-SJBS-202501), Knowledge Innovation Program of Chinese Academy of Sciences (Institute of Deep-sea Science and Engineering, Chinese Academy of Sciences; grant number Y970011001), Hainan Provincial Funding for Outstanding Talent Team Development (Hainan Provincial Talent Development Bureau; grant number E2500001) to M.C.Y.L.V.

## Notes

### Competing Interest Statement

The authors have declared no competing interest.

