## Supplementary materials (figures and tables) for "Biosurfactant production under cold-saline stress is a strain specific trait independent from phylogeny and biosynthetic gene content"

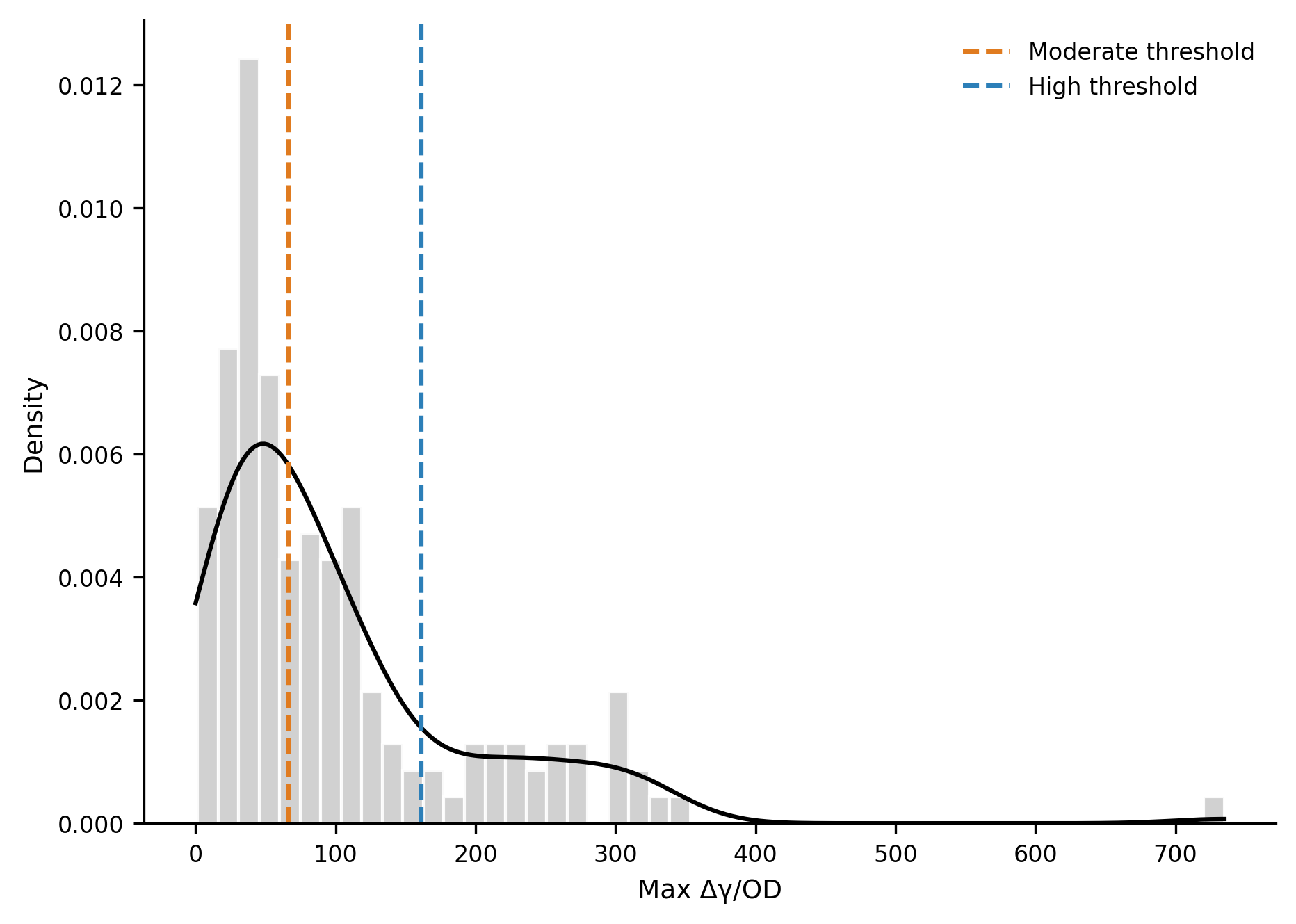

**Figure S1. Data-driven producer classification by Gaussian mixture modelling.** Distribution of maximum Δγ/OD values pooled across all strains and conditions (grey histogram) with the fitted kernel density (black line). Dashed lines mark the component boundaries identified by a Gaussian finite mixture model (Bayesian Information Criterion-selected), yielding data-driven thresholds separating Low, Moderate (orange), and High (blue) producer classes.

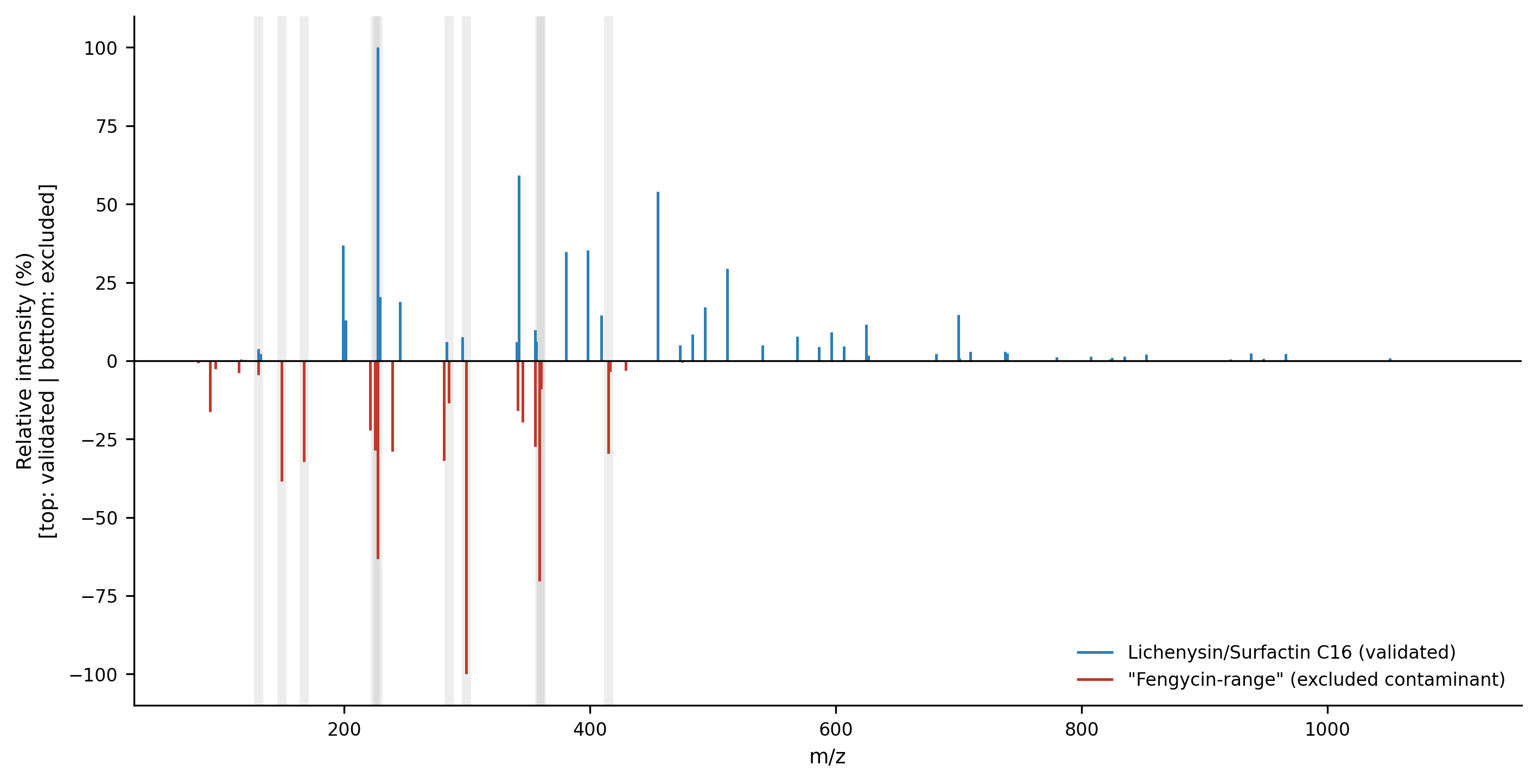

**Figure S2. MS² validation distinguishing a genuine lipopeptide from a contaminant-dominated cluster.** Mirror plot comparing the MS² spectrum of the validated Lichenysin/Surfactin C16 homolog (cluster 128273; top, blue) with that of an excluded "Fengycin-range" cluster (cluster 112611; bottom, red). Grey vertical bands mark fragment ions shared between clusters of different precursor mass, a non-peptidic fingerprint diagnostic of a chemical contaminant, confirming the exclusion of the Fengycin-range features from biological interpretation.

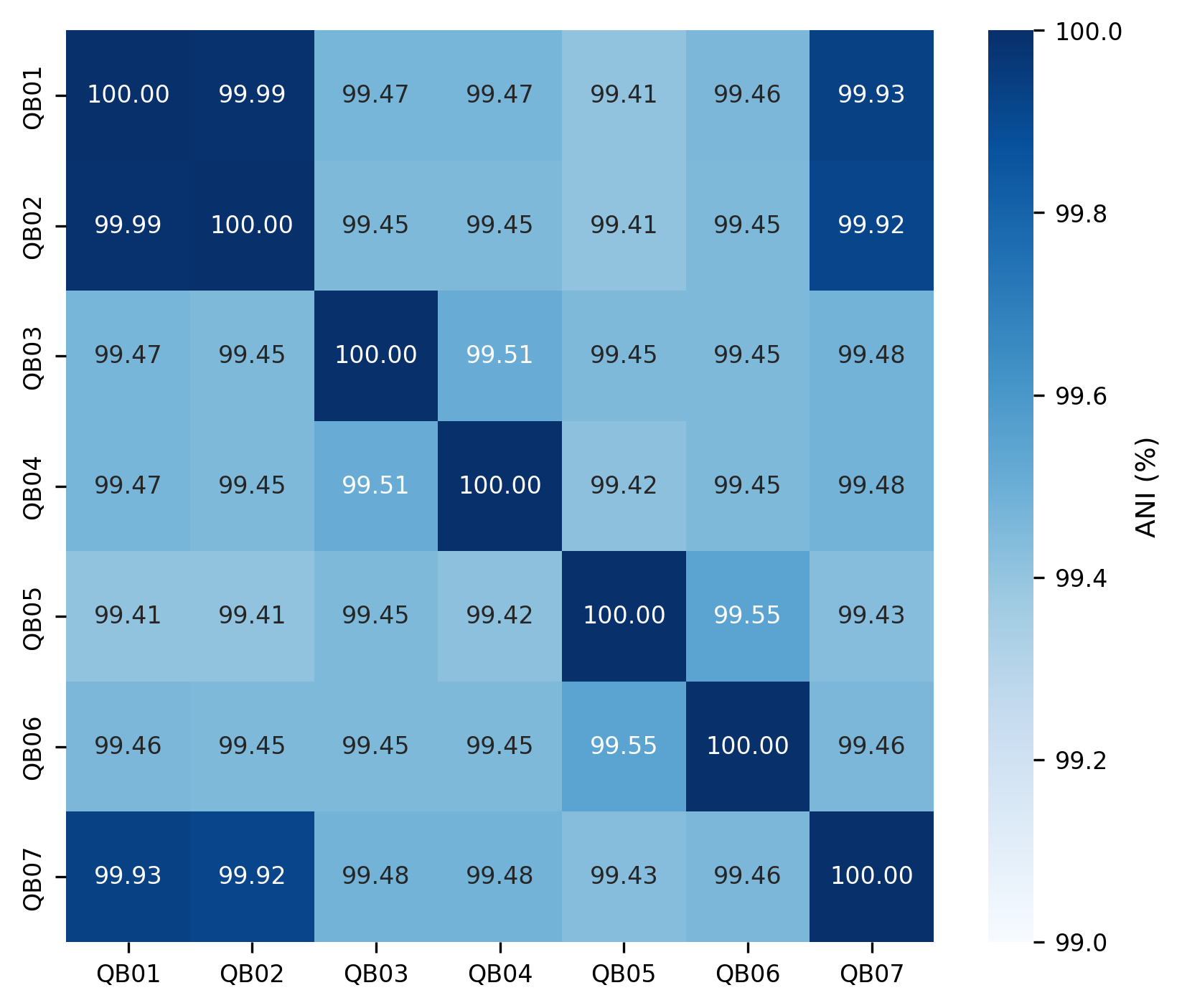

**Figure S3. Pairwise average nucleotide identity within the *B. zhangzhouensis* clade.** Heatmap of pairwise ANI (%) among the seven sequenced clade members (QB01–QB07), computed with FastANI. All values exceed 99.4%, confirming near-clonal relatedness; QB01 and QB02 share 99.99% ANI, consistent with effectively identical genomes.

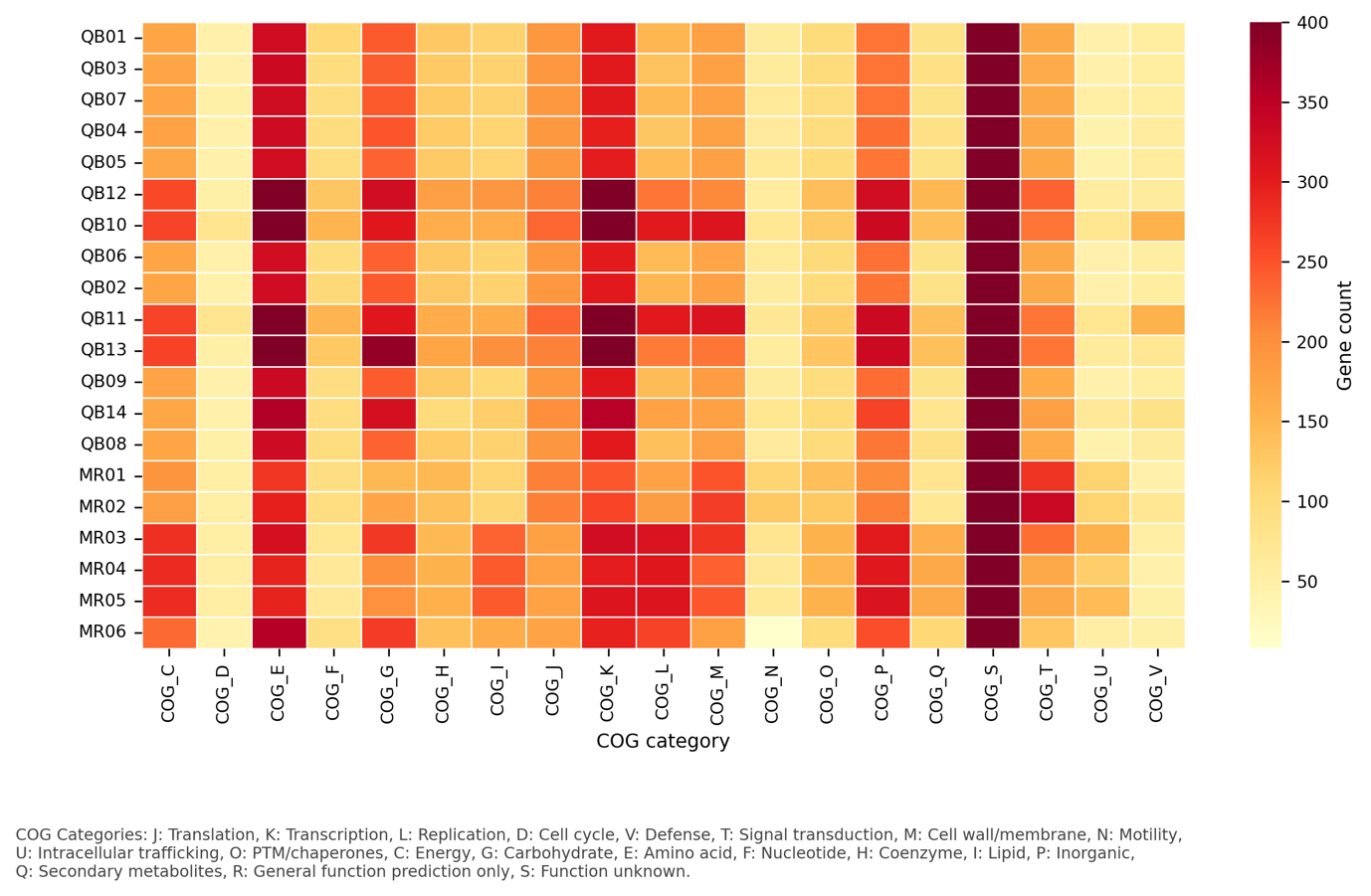

**Figure S4. Functional gene content across the collection by COG category.** Heatmap of gene counts per Cluster of Orthologous Genes (COG) functional category (columns) for each sequenced genome (rows), annotated by eggNOG-mapper. The colour scale is capped at 400 genes to preserve contrast across categories; COG single-letter codes are defined in the footnote. Strain identifiers follow Table 1.

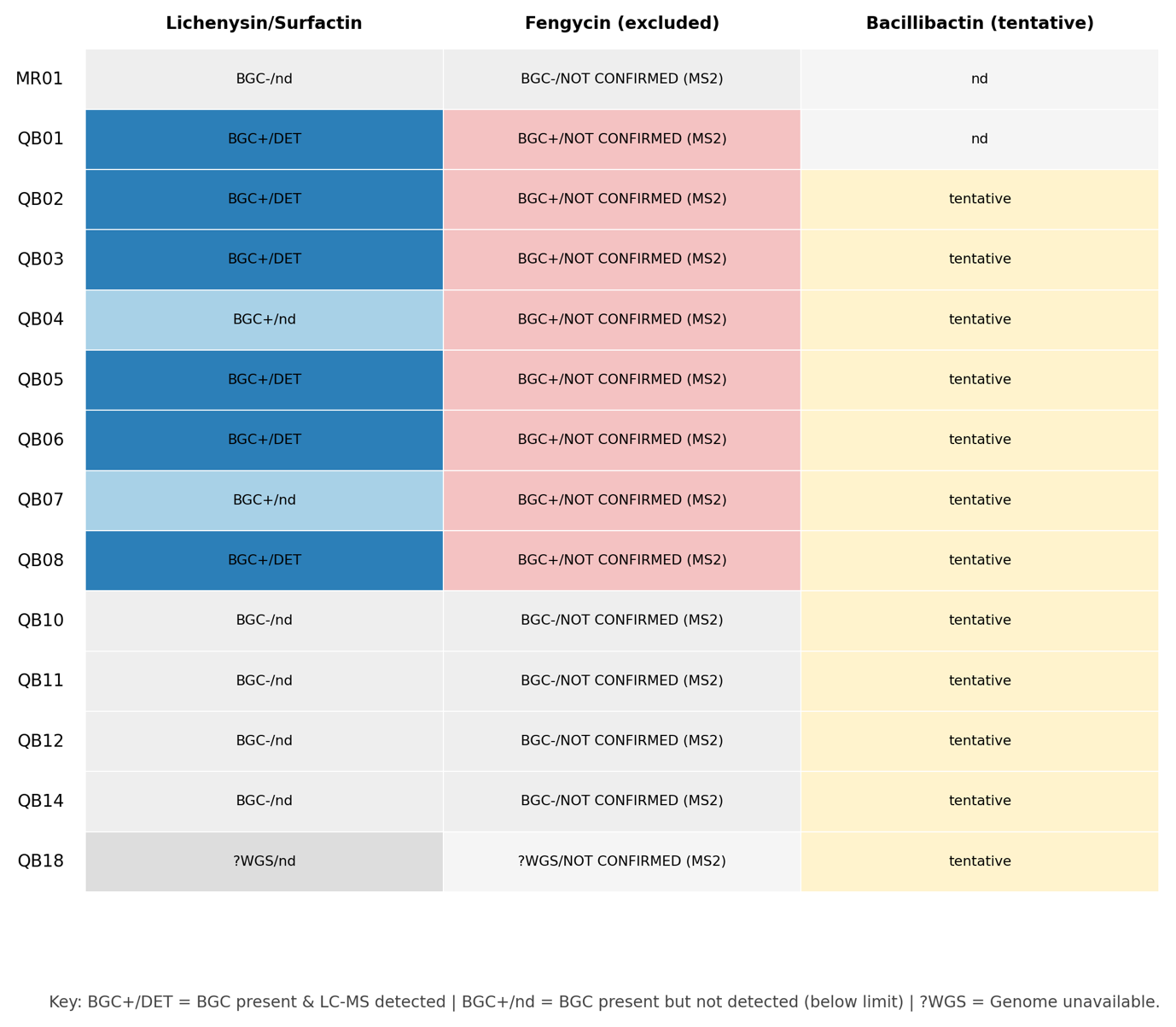

**Figure S5. Concordance between genomically predicted and LC-MS-detected biosurfactants.** Matrix summarising, for each LC-MS-analysed strain (rows), the genomic prediction and chemical detection status of three biosurfactant classes (columns): Lichenysin/Surfactin, Fengycin, and Bacillibactin. Cell labels indicate BGC presence and detection outcome (BGC+/DET, cluster present and compound detected; BGC+/nd, cluster present but not detected; BGC−/nd, cluster absent and not detected). Fengycin was genomically predicted but not confirmed by MS² and is therefore reported as excluded; Bacillibactin is reported as a tentative annotation. Strain identifiers follow Table 1.

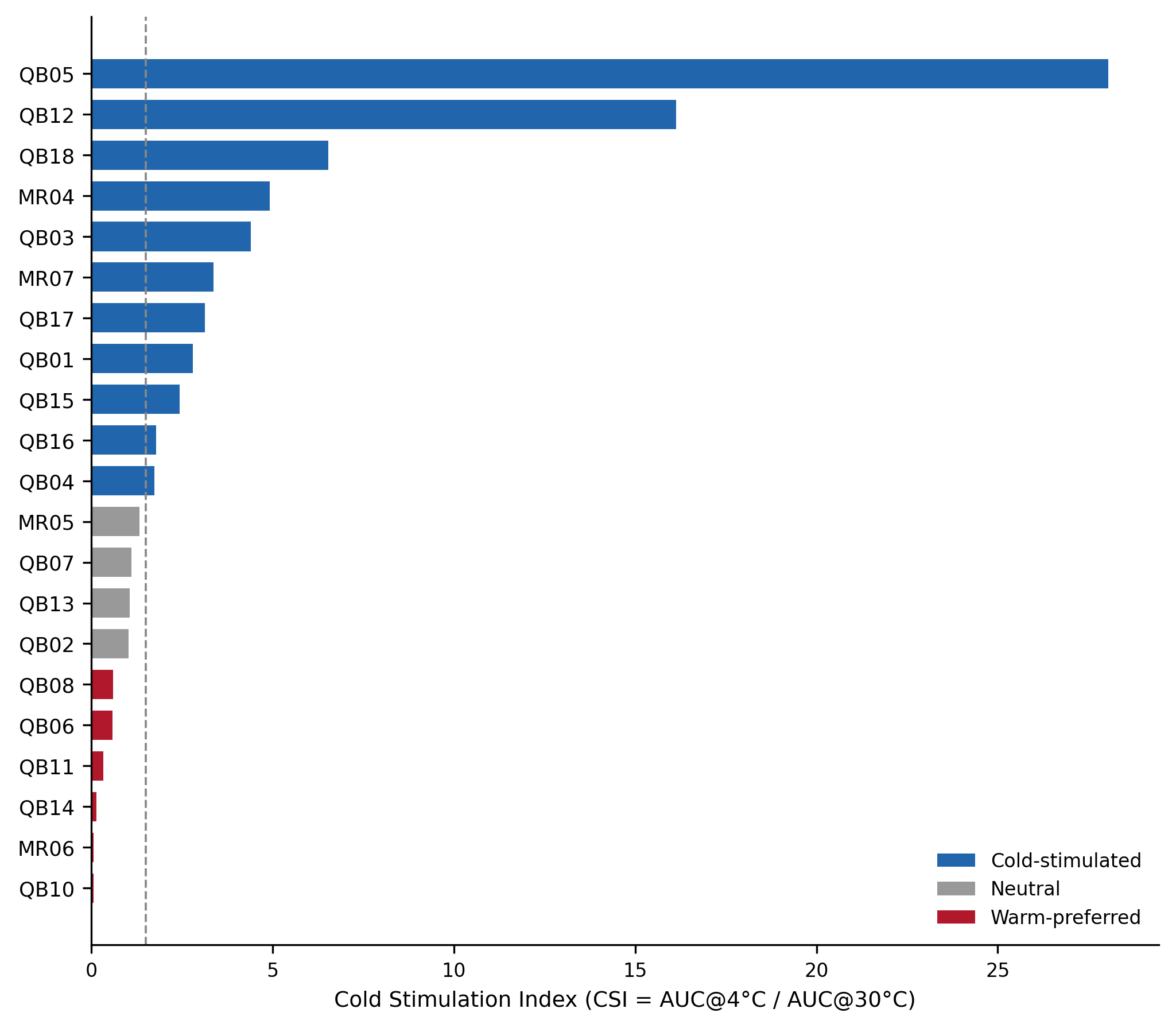

**Figure S6. Per-strain cold-stimulation index ranking.** Cold-stimulation index (CSI = AUC(Δγ/OD) at 4 °C ÷ AUC(Δγ/OD) at 30 °C) for all 22 phenotyped strains, ranked in descending order and coloured by cold-response class (cold-stimulated, blue; neutral, grey; warm-preferred, red). The dashed line marks the cold-stimulation threshold (CSI = 1.5). Strain identifiers follow Table 1.

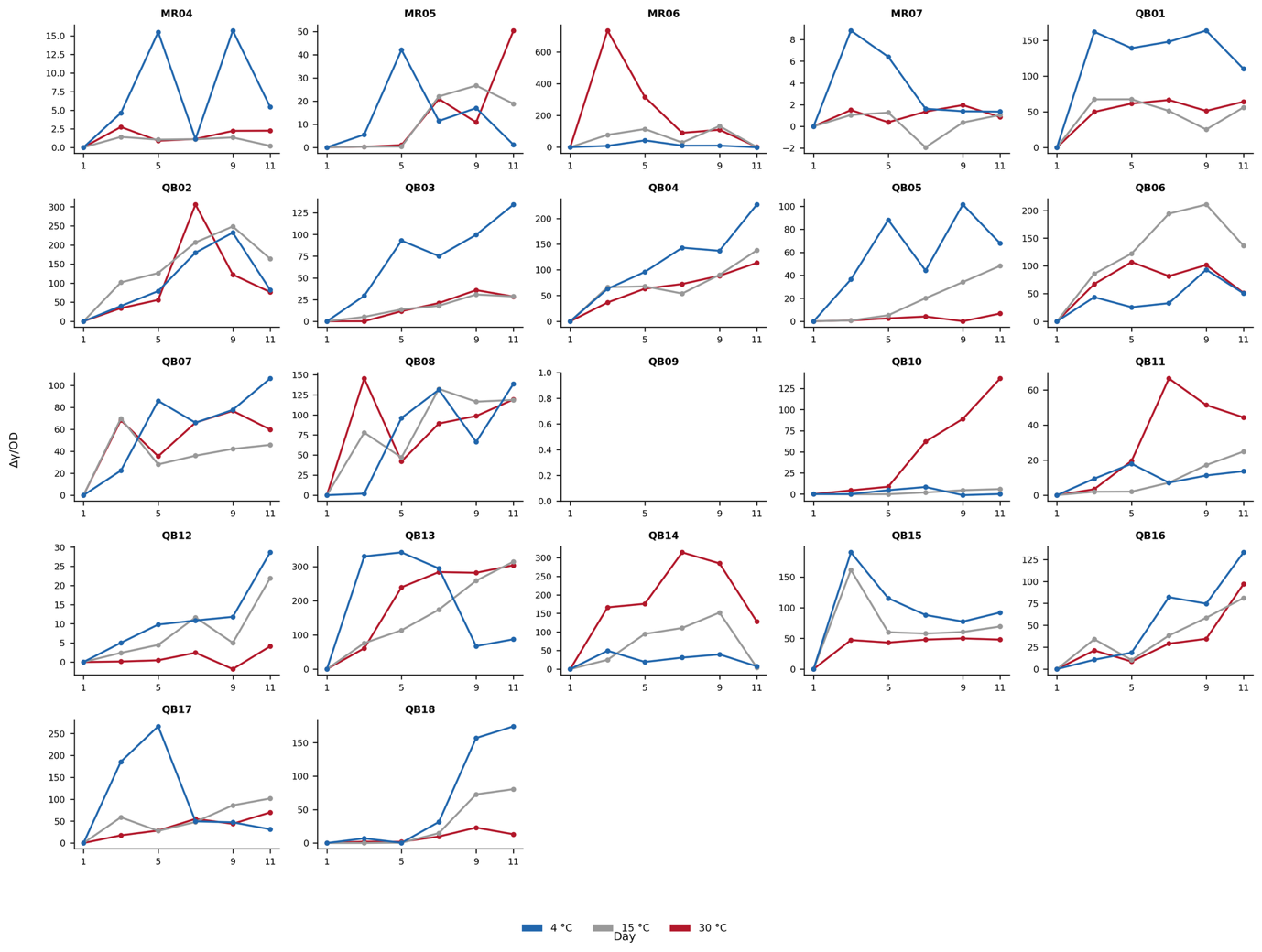

**Figure S7. Biosurfactant activity time-series for all strains under temperature-only conditions.** Mean Δγ/OD over the 11-day incubation for each of the 22 phenotyped strains (individual panels), at three temperatures (4 °C, blue; 15 °C, grey; 30 °C, red). Per-strain y-axis scaling reflects the wide range of production magnitudes across the collection. Strain identifiers follow Table 1.

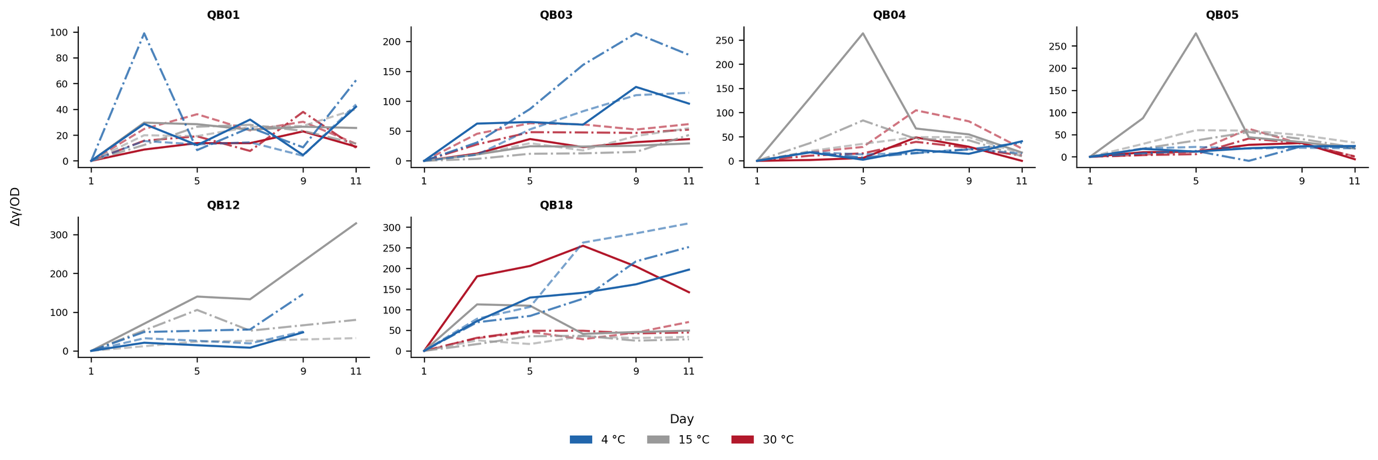

**Figure S8. Biosurfactant activity time-series under combined temperature × salinity stress.** Mean Δγ/OD over the 11-day incubation for the cold-stimulated strains tested in the full factorial experiment (individual panels). Line colour denotes temperature (4 °C, blue; 15 °C, grey; 30 °C, red) and line style denotes NaCl concentration (solid, 1.5 M; dash-dot, 1.0 M; dashed, 0.5 M; dotted, 0 M). Strain identifiers follow Table 1.

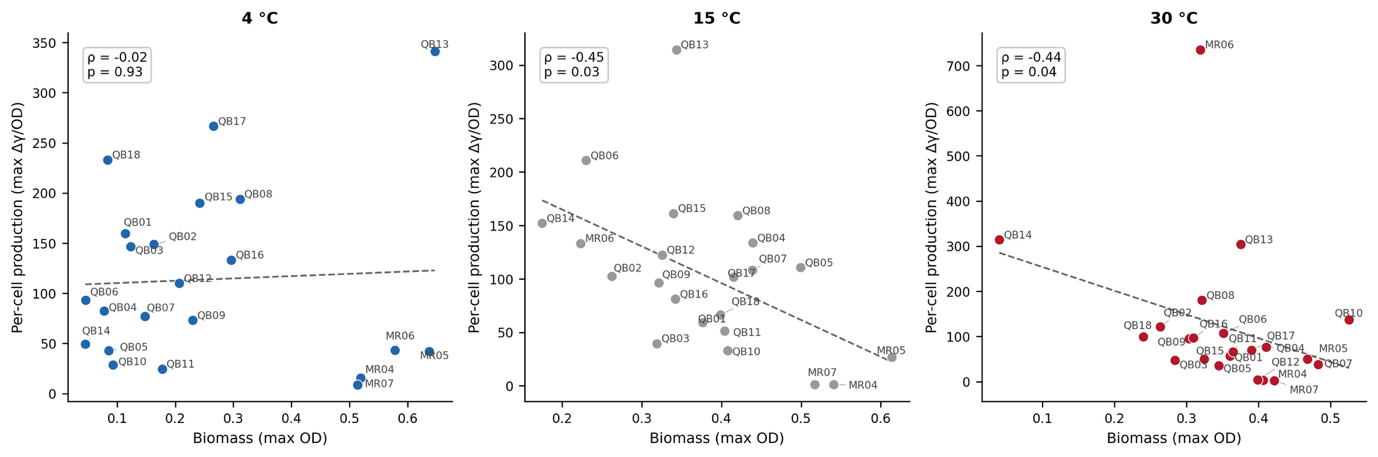

**Figure S9. Biosurfactant production is decoupled from growth at each temperature.** Per-cell production (maximum Δγ/OD) plotted against biomass (maximum OD) for all phenotyped strains at 4 °C, 15 °C, and 30 °C (separate panels), with Spearman correlations annotated (4 °C, ρ = −0.02, *p* = 0.93; 15 °C, ρ = −0.49, *p* = 0.03; 30 °C, ρ = −0.41, *p* = 0.04). Dashed lines show linear fits. The negative or null associations indicate that production is not biomass-driven. Strain identifiers follow Table 1.

Table S1. Analytical methods, software, and key parameters.

| **Parameter** | **Value / description** |
| --- | --- |
| LC-MS instrument | Thermo Orbitrap, H-ESI positive mode |
| Data conversion | .raw to .mzML via ThermoRawFileParser |
| Molecular networking | GNPS METABOLOMICS-SNETS-V2 (job ID: 620d65e3) |
| Spectral features | 5,726 features across 522 spectral families |
| LC-MS samples | 14 samples |
| Precursor ion mass tolerance | 2.0 Da |
| Fragment ion mass tolerance | 0.5 Da |
| Dereplication | DEREPLICATOR+ (lipopeptide structural confirmation) |
| Genome assembly quality | All 20 genomes: 100% complete, <=2.2% contamination (CheckM) |
| Taxonomy assignment | GTDB-Tk (Genome Taxonomy Database reference) |
| BGC annotation | antiSMASH v7 |
| Pan-genome analysis | Panaroo (9 Bacillus genomes; 4,625 gene families, 68.9% core) |
| Core-genome phylogeny | IQ-TREE2 (ML, core-genome alignment from Panaroo) |
| Physiology experiments | Surface tension / OD; 22 strains x 3 temperatures x 4 salinities; 6 timepoints (day 1-11) |
| CSI definition | AUC(dg/OD)at_4C / AUC(dg/OD)at_30C (temperature-only experiments) |
| Cold response classification | Cold-stimulated (CSI > 1.5), Neutral (0.7-1.5), Warm-preferred (< 0.7) |
| Statistical software | R 4.x (lme4, emmeans, mclust); Python 3.11 (scipy, scikit-bio, pandas) |

Table S2. Statistical test results for all key comparisons.

| **Test / comparison** | **Method** | **Statistic** | **p-value** | **Effect size / detail** | **n** | **Interpretation** |
| --- | --- | --- | --- | --- | --- | --- |
| Temperature effect (community) | LMM ANOVA | F(2,45.9) = 1.84 | 0.17 | R2m = 0.018, R2c = 0.68 | 168 | ns at community level; strain identity dominates |
| Temp x salinity interaction | LMM ANOVA | F(4,70.3) = 1.86 | 0.13 | — | 168 | ns; response is strain-specific |
| Cold-salt synergy | Bliss independence | 8/8 synergistic | — | all synergy indices > 0 | 8 | All tested strains synergistic |
| Growth-production decoupling @4C | Spearman | rho = -0.35 | 0.11 | — | 23 | ns (production independent of growth at 4C) |
| Growth-production decoupling @15C | Spearman | rho = -0.75 | < 0.001 | — | 23 | Significant negative correlation |
| Growth-production decoupling @30C | Spearman | rho = -0.41 | 0.05 | — | 23 | Borderline significant |
| Regulatory genes vs production @4C | Spearman | rho = 0.85 | < 0.001 | — | 17 | Strong positive (regulatory dosage predicts production) |
| Lipopeptide BGC presence vs production @4C | Wilcoxon rank-sum | W = 16 | 0.06 | r = 0.56; median 194 vs 43 | 17 | BGC+ higher; borderline significant |
| Total BGC count vs production @4C | Spearman | rho = -0.06 | 0.80 | — | 17 | Null (gene number not predictive) |
| Phylogenetic signal in cold trait | Mantel (permutation) | r = 0.14 | 0.28 | — | 7 | No phylogenetic signal within clade |
| BGC density Qaidam vs Marine | t-test | 2.80 vs 1.45 BGC/Mb | < 0.01 | — | 20 | Qaidam enriched in BGCs |
| Cold production Qaidam vs Marine @4C | Mann-Whitney | U = 38 | 0.03 | median 156.0 vs 42.0 | 17 | Qaidam produces more under cold stress |
| COG functional profiles (PCoA) | PERMANOVA | R2 = 0.67 | 0.001 | — | 20 | Environment structures functional gene content |
| COG environment effect | PERMANOVA | R2 = 0.58 | 0.001 | — | 20 | Environment dominates functional variation |

Table S3. Biosynthetic gene cluster (BGC) presence/absence across 20 sequenced genomes.

| **Strain ID** | **Lichenysin** | **Surfactin** | **Fengycin** | **Bacillibactin** | **Bacilysin** | **Zwittermicin** | **Desmamide** | **Koranimine** | **Subtilomycin** | **Arylpolyene** | **Ectoine** | **Zeaxanthin** | **Lassopeptide** | **T3PKS** | **NAPAA** | **RiPP** | **Terpene** |
| --- | --- | --- | --- | --- | --- | --- | --- | --- | --- | --- | --- | --- | --- | --- | --- | --- | --- |
| MR01 | 0 | 0 | 0 | 0 | 0 | 0 | 0 | 0 | 0 | 1 | 0 | 0 | 0 | 0 | 0 | 1 | 0 |
| MR02 | 0 | 0 | 0 | 0 | 0 | 0 | 0 | 0 | 0 | 1 | 0 | 0 | 0 | 0 | 0 | 1 | 0 |
| MR03 | 0 | 0 | 0 | 0 | 0 | 0 | 0 | 0 | 0 | 0 | 1 | 0 | 1 | 0 | 0 | 0 | 1 |
| MR04 | 0 | 0 | 0 | 0 | 0 | 0 | 0 | 0 | 0 | 0 | 1 | 1 | 1 | 0 | 0 | 0 | 0 |
| MR05 | 0 | 0 | 0 | 0 | 0 | 0 | 0 | 0 | 0 | 0 | 1 | 1 | 1 | 0 | 0 | 0 | 1 |
| MR06 | 0 | 0 | 0 | 0 | 0 | 0 | 0 | 0 | 0 | 0 | 0 | 0 | 0 | 1 | 1 | 1 | 1 |
| QB01 | 1 | 0 | 1 | 1 | 1 | 1 | 0 | 0 | 0 | 0 | 0 | 0 | 0 | 0 | 0 | 1 | 1 |
| QB02 | 1 | 0 | 1 | 1 | 1 | 1 | 0 | 0 | 0 | 0 | 0 | 0 | 0 | 0 | 0 | 1 | 1 |
| QB03 | 0 | 1 | 1 | 1 | 1 | 0 | 0 | 0 | 0 | 0 | 0 | 0 | 0 | 0 | 0 | 1 | 1 |
| QB04 | 1 | 0 | 1 | 1 | 1 | 0 | 0 | 0 | 0 | 0 | 0 | 0 | 0 | 0 | 0 | 1 | 1 |
| QB05 | 1 | 0 | 1 | 1 | 1 | 0 | 0 | 0 | 0 | 0 | 0 | 0 | 0 | 0 | 0 | 1 | 1 |
| QB06 | 1 | 0 | 1 | 1 | 1 | 1 | 0 | 0 | 0 | 0 | 0 | 0 | 0 | 0 | 0 | 1 | 1 |
| QB07 | 1 | 0 | 1 | 1 | 1 | 1 | 0 | 0 | 0 | 0 | 0 | 0 | 0 | 0 | 0 | 1 | 1 |
| QB08 | 1 | 0 | 1 | 1 | 1 | 0 | 0 | 0 | 0 | 0 | 0 | 0 | 0 | 0 | 0 | 1 | 1 |
| QB09 | 1 | 0 | 1 | 1 | 1 | 0 | 0 | 0 | 0 | 0 | 0 | 0 | 0 | 0 | 0 | 1 | 1 |
| QB10 | 0 | 0 | 1 | 1 | 0 | 1 | 0 | 0 | 1 | 0 | 0 | 0 | 0 | 0 | 0 | 1 | 1 |
| QB11 | 0 | 0 | 1 | 1 | 0 | 1 | 0 | 0 | 1 | 0 | 0 | 0 | 0 | 0 | 0 | 1 | 1 |
| QB12 | 0 | 0 | 1 | 0 | 0 | 0 | 1 | 1 | 0 | 0 | 0 | 0 | 0 | 0 | 0 | 0 | 1 |
| QB13 | 0 | 0 | 1 | 0 | 0 | 0 | 0 | 1 | 0 | 0 | 0 | 0 | 0 | 0 | 0 | 1 | 1 |
| QB14 | 0 | 0 | 0 | 0 | 0 | 0 | 0 | 0 | 0 | 0 | 1 | 0 | 0 | 1 | 0 | 0 | 1 |

1 = present, 0 = absent. Annotated by antiSMASH v7.

Table S4. Per-strain physiological metrics, temperature-only experiments (22 strains).

| **Strain ID** | **Max dg/OD (ever)** | **Mean dg/OD** | **Max dg/OD @4C** | **Max dg/OD @30C** | **AUC @4C** | **AUC @30C** | **CSI** | **Cold response class** | **Producer class** |
| --- | --- | --- | --- | --- | --- | --- | --- | --- | --- |
| MR04 | 15.68 | 6.61 | 15.68 | 2.74 | 79.23 | 16.13 | 4.91 | Cold-stimulated | Low |
| MR05 | 50.35 | 39.68 | 42.04 | 50.35 | 152.87 | 116.18 | 1.32 | Neutral | Low |
| MR06 | 735 | 303.79 | 43.13 | 735 | 142.38 | 2505.01 | 0.06 | Warm-preferred | High |
| MR07 | 8.82 | 4.02 | 8.82 | 1.96 | 37.88 | 11.28 | 3.36 | Cold-stimulated | Low |
| QB01 | 308.47 | 91.84 | 308.47 | 105.44 | 1661.53 | 593.24 | 2.8 | Cold-stimulated | High |
| QB02 | 305.94 | 124.17 | 232.17 | 305.94 | 1144.53 | 1113.82 | 1.03 | Neutral | High |
| QB03 | 213.53 | 77.66 | 213.53 | 63.06 | 728.48 | 165.53 | 4.4 | Cold-stimulated | High |
| QB04 | 263.94 | 97.55 | 227.04 | 113.63 | 1105.89 | 635.43 | 1.74 | Cold-stimulated | High |
| QB05 | 278.35 | 63.06 | 101.55 | 63.14 | 608.34 | 21.69 | 28.05 | Cold-stimulated | High |
| QB06 | 210.8 | 136.99 | 93.18 | 107 | 440.04 | 765.97 | 0.57 | Warm-preferred | High |
| QB07 | 240.48 | 74.44 | 106.31 | 76.79 | 610.34 | 553.07 | 1.1 | Neutral | High |
| QB08 | 274.71 | 177.74 | 194.21 | 274.71 | 773.52 | 1292.86 | 0.6 | Warm-preferred | High |
| QB09 | 131.19 | 88.12 | 117.77 | 120.49 |  |  |  |  | Moderate |
| QB10 | 136.91 | 42.55 | 38.82 | 136.91 | 23.62 | 465.38 | 0.05 | Warm-preferred | Moderate |
| QB11 | 75.68 | 41.06 | 34.82 | 66.51 | 105.08 | 326 | 0.32 | Warm-preferred | Moderate |
| QB12 | 329.12 | 103.69 | 229.88 | 4.12 | 103.64 | 6.42 | 16.13 | Cold-stimulated | High |
| QB13 | 341.37 | 319.83 | 341.37 | 303.77 | 2153.18 | 2035.42 | 1.06 | Neutral | High |
| QB14 | 314.25 | 171.92 | 49.41 | 314.25 | 285.45 | 2009.7 | 0.14 | Warm-preferred | High |
| QB15 | 189.86 | 133.7 | 189.86 | 49.98 | 1033.15 | 425.04 | 2.43 | Cold-stimulated | High |
| QB16 | 133.09 | 103.77 | 133.09 | 97.07 | 505.32 | 284.2 | 1.78 | Cold-stimulated | Moderate |
| QB17 | 266.58 | 146.12 | 266.58 | 69.97 | 1129.84 | 360.32 | 3.14 | Cold-stimulated | High |
| QB18 | 308.96 | 132.82 | 308.96 | 254.66 | 566.17 | 86.72 | 6.53 | Cold-stimulated | High |

dg/OD = surface tension reduction normalized to optical density. AUC = area under the time-series curve. CSI from temperature-only experiments.

**Table S5. Per-condition physiological metrics, combined temperature x salinity experiments (11 strains x 6 conditions).**

| **Strain ID** | **Temperature** | **NaCl (M)** | **Max dg/OD** | **AUC dg/OD** | **Max OD** |
| --- | --- | --- | --- | --- | --- |
| QB01 | 4C | 0.5 | 43.8 | 136.9 | 0.209 |
| QB01 | 4C | 1 | 99 | 349.9 | 0.13 |
| QB01 | 4C | 1.5 | 42 | 196.9 | 0.045 |
| QB01 | 15C | 0.5 | 40.1 | 225.2 | 0.388 |
| QB01 | 15C | 1 | 28 | 193.6 | 0.383 |
| QB01 | 15C | 1.5 | 29.6 | 243 | 0.453 |
| QB01 | 30C | 0.5 | 36.2 | 243.1 | 0.322 |
| QB01 | 30C | 1 | 38 | 170.9 | 0.426 |
| QB01 | 30C | 1.5 | 22.9 | 129.4 | 0.431 |
| QB02 | 4C | 0.5 | 99.6 | 420.2 | 0.213 |
| QB02 | 4C | 1 | 93.6 | 604.4 | 0.205 |
| QB02 | 4C | 1.5 | 169.6 | 864.3 | 0.185 |
| QB02 | 15C | 0.5 | 97.8 | 601.5 | 0.211 |
| QB02 | 15C | 1 | 38.1 | 204.6 | 0.228 |
| QB02 | 15C | 1.5 | 25.2 | 114.4 | 0.347 |
| QB02 | 30C | 0.5 | 87.4 | 518.6 | 0.261 |
| QB02 | 30C | 1 | 42.6 | 250.5 | 0.28 |
| QB02 | 30C | 1.5 | 49.6 | 206.4 | 0.238 |
| QB03 | 4C | 0.5 | 113.9 | 626.5 | 0.18 |
| QB03 | 4C | 1 | 213.5 | 1161.1 | 0.086 |
| QB03 | 4C | 1.5 | 123.8 | 719.5 | 0.06 |
| QB03 | 15C | 0.5 | 54.7 | 253 | 0.235 |
| QB03 | 15C | 1 | 43.3 | 129.1 | 0.289 |
| QB03 | 15C | 1.5 | 29.2 | 198.5 | 0.358 |
| QB03 | 30C | 0.5 | 63.1 | 505.4 | 0.295 |
| QB03 | 30C | 1 | 52.4 | 392.4 | 0.266 |
| QB03 | 30C | 1.5 | 36.6 | 242.7 | 0.272 |
| QB04 | 4C | 0.5 | 24.3 | 160.6 | 0.089 |
| QB04 | 4C | 1 | 37.6 | 165.5 | 0.048 |
| QB04 | 4C | 1.5 | 40.6 | 155.4 | 0.038 |
| QB04 | 15C | 0.5 | 49.3 | 322.7 | 0.455 |
| QB04 | 15C | 1 | 83.7 | 426.6 | 0.572 |
| QB04 | 15C | 1.5 | 263.9 | 1049.8 | 0.329 |
| QB04 | 30C | 0.5 | 104.6 | 492.1 | 0.346 |
| QB04 | 30C | 1 | 39.4 | 196.6 | 0.384 |
| QB04 | 30C | 1.5 | 48.6 | 171 | 0.522 |
| QB05 | 4C | 0.5 | 22.3 | 179.3 | 0.075 |
| QB05 | 4C | 1 | 23.9 | 93.3 | 0.07 |
| QB05 | 4C | 1.5 | 24.3 | 169.5 | 0.042 |
| QB05 | 15C | 0.5 | 60 | 425.6 | 0.407 |
| QB05 | 15C | 1 | 55.8 | 323.6 | 0.522 |
| QB05 | 15C | 1.5 | 278.3 | 903.9 | 0.511 |
| QB05 | 30C | 0.5 | 63.1 | 217.4 | 0.362 |
| QB05 | 30C | 1 | 41.2 | 164 | 0.394 |
| QB05 | 30C | 1.5 | 31.2 | 155 | 0.271 |
| QB07 | 4C | 0.5 | 48 | 160.1 | 0.161 |
| QB07 | 4C | 1 | 102.2 | 264 | 0.058 |
| QB07 | 4C | 1.5 | 52 | 315.4 | 0.043 |
| QB07 | 15C | 0.5 | 68.4 | 348.2 | 0.421 |
| QB07 | 15C | 1 | 53.8 | 357 | 0.429 |
| QB07 | 15C | 1.5 | 240.5 | 671.7 | 0.323 |
| QB07 | 30C | 0.5 | 15.5 | 93.1 | 0.637 |
| QB07 | 30C | 1 | 34.4 | 151.7 | 0.423 |
| QB07 | 30C | 1.5 | 25.6 | 95.2 | 0.544 |
| QB09 | 4C | 0.5 | 36.8 | 229.7 | 0.213 |
| QB09 | 4C | 1 | 117.8 | 557.8 | 0.202 |
| QB09 | 4C | 1.5 | 64.6 | 341.2 | 0.275 |
| QB09 | 15C | 0.5 | 131.2 | 821.8 | 0.229 |
| QB09 | 15C | 1 | 87.9 | 598.2 | 0.346 |
| QB09 | 15C | 1.5 | 69.6 | 379 | 0.388 |
| QB09 | 30C | 0.5 | 120.5 | 840.4 | 0.308 |
| QB09 | 30C | 1 | 110.8 | 737.9 | 0.278 |
| QB09 | 30C | 1.5 | 53.8 | 348.1 | 0.324 |
| QB10 | 4C | 0.5 | 38.8 | 194.9 | 0.046 |
| QB10 | 4C | 1 | 29.1 | 147.8 | 0.048 |
| QB10 | 4C | 1.5 | 37.6 | 210.2 | 0.047 |
| QB10 | 15C | 0.5 | 32.6 | 130.2 | 0.403 |
| QB10 | 15C | 1 | 43.1 | 324.8 | 0.302 |
| QB10 | 15C | 1.5 | 50.5 | 363.2 | 0.292 |
| QB10 | 30C | 0.5 | N.D | N.D | 0.433 |
| QB10 | 30C | 1 | N.D | N.D | 0.293 |
| QB10 | 30C | 1.5 | N.D | N.D | 0.252 |
| QB11 | 4C | 0.5 | 34.8 | 139.8 | 0.047 |
| QB11 | 4C | 1 | 30.5 | 178.8 | 0.058 |
| QB11 | 4C | 1.5 | 14.9 | 92.9 | 0.053 |
| QB11 | 15C | 0.5 | 31.6 | 115.8 | 0.465 |
| QB11 | 15C | 1 | 72.8 | 376.5 | 0.272 |
| QB11 | 15C | 1.5 | 75.7 | 438.3 | 0.175 |
| QB11 | 30C | 0.5 | N.D | N.D | 0.296 |
| QB11 | 30C | 1 | N.D | N.D | 0.173 |
| QB11 | 30C | 1.5 | N.D | N.D | 0.184 |
| QB12 | 4C | 0.5 | 97.3 | 206.3 | 0.138 |
| QB12 | 4C | 1 | 229.9 | 458.3 | 0.054 |
| QB12 | 4C | 1.5 | 83.7 | 135.8 | 0.064 |
| QB12 | 15C | 0.5 | 32.9 | 216.3 | 0.396 |
| QB12 | 15C | 1 | 105.6 | 633.3 | 0.26 |
| QB12 | 15C | 1.5 | 329.1 | 1478.5 | 0.046 |
| QB12 | 30C | 0.5 | N.D | N.D | 0.399 |
| QB12 | 30C | 1 | N.D | N.D | 0.217 |
| QB12 | 30C | 1.5 | N.D | N.D | 0.053 |
| QB18 | 4C | 0.5 | 309 | 1771.5 | 0.048 |
| QB18 | 4C | 1 | 251.6 | 1246.1 | 0.046 |
| QB18 | 4C | 1.5 | 196.8 | 1203.6 | 0.047 |
| QB18 | 15C | 0.5 | 35.7 | 251.8 | 0.419 |
| QB18 | 15C | 1 | 36.6 | 254.4 | 0.379 |
| QB18 | 15C | 1.5 | 112.6 | 666.9 | 0.247 |
| QB18 | 30C | 0.5 | 70 | 369.3 | 0.248 |
| QB18 | 30C | 1 | 48.8 | 387.8 | 0.284 |
| QB18 | 30C | 1.5 | 254.7 | 1832.7 | 0.068 |

dg/OD = surface tension reduction normalized to OD. N.D = Not Detected (no measurable surface tension reduction). 11 strains tested at 3 temperatures x 3 NaCl concentrations.
